# Streamlined DNA template preparation and co-transcriptional 5′ capped RNA synthesis enabled by solid-phase catalysis

**DOI:** 10.1101/2023.10.28.564520

**Authors:** Guillermo García-Marquina, Aihua Zhang, Michael Sproviero, Yi Fang, Andrew F. Gardner, G. Brett Robb, S. Hong Chan, Ming-Qun Xu

## Abstract

The success of SARS-CoV-2 mRNA vaccines demonstrated that rapid, large-scale manufacturing of synthetic mRNA is necessary for an effective and timely response to a pandemic. Innovations in areas such as template design and manufacturing processes are being implemented to facilitate more simple, cost-effective and scalable mRNA synthesis. In this study, for the first time, we demonstrate that the enzymatic steps in mRNA production (including DNA template linearization, RNA synthesis, 5′ capping and methylation) can be carried out using enzymes immobilized to a solid support. Specifically, we demonstrate efficient IVT template DNA linearization using immobilized BspQI, where the linearized template DNA can be directly used in IVT without the need of purification. We also showed that immobilized T7 RNA polymerase, Faustovirus RNA capping enzyme (FCE), vaccinia cap 2′-O-methyltransfease (2′OMTase) and a novel FCE::T7RNAP fusion enable efficient enzymatic synthesis of Cap-1 RNA in a one-pot format. This solid-phase enzymatic platform may enable highly efficient, seamless and continuous mRNA synthesis workflows that minimizes sample loss and units of operation in biopharmaceutical manufacturing.

## MAIN

Protein expression driven by exogenous synthetic mRNA was first reported in mice in 1990.^1^ Since then, research and development towards realizing synthetic mRNA therapeutics progressed steadily until 2020, when synthetic mRNA was used during the SARS-CoV-2 pandemic as a vaccine.^2^ mRNA therapeutics have rapidly expanded since then, with many reaching clinical trials for treatment of various diseases, including heart failure and a congenital liver-specific storage disease.^3^ Without the need to enter the nucleus, mRNA is a more effective and safer vaccine and therapeutic modality than DNA because of the faster protein expression, minimal risk of genome incorporation and the natural cellular degradation processes.^4, 5^ Moreover, *in vitro* transcription (IVT) is a cell-free production process that allows for tight control of reagents and undesirable biological contaminants. As a nucleic acid, the functionality of synthetic mRNA largely derives from its nucleotide sequence with little variation in chemical properties. Therefore, manufacturing of multiple synthetic mRNA drug substances can be readily standardized compared to recombinant proteins or small molecules.^6, 7^ Thus far, DNA-dependent RNA polymerase (RNAP) derived from bacteriophage T7 is commonly used in IVT because of its simplicity (single protein with well-defined short promoter sequences) and high transcription efficiency.^8–11^

A functional IVT mRNA contains unique modules that contribute to the function and regulation of the molecule *in vivo*. Similar to endogenous mRNA, the 5′ end is modified with a 7-methylguanosine nucleoside linked through a triphosphate group to the 5′ end of mRNA. In mammals, the first nucleotide at the 5′ end is methylated at the 2′-O position of the ribose. This cap structure (Cap-1) contributes to efficient translation and evading innate immune response, among other functions. Proceeding the cap structure is a 5′ untranslated region (UTR), the open reading frame encoding the target protein, and 3′ UTR, followed by a poly(A) tail at the 3′ end. The poly(A) binding proteins and translation initiation factor proteins circularize the mRNA and recruit ribosomes for initiating translation, as well as regulating the lifespan of the mRNA molecules.^12, 13^

*In vitro* mRNA synthesis involves multiple steps: DNA template preparation (comprised of plasmid preparation and linearization), the IVT reaction with templated poly(A) tail, 5′ capping and poly(A) tail addition if post-IVT enzymatic poly(A) tailing is desired. While plasmid preparation is typically done using *Escherichia coli* as the host for propagation followed by plasmid isolation,^14–16^ plasmid linearization, IVT and post-transcriptional capping involves multiple enzymes.^16–21^ Current workflows use separate enzymatic reactions for each of the steps. Purification of the product from the respective enzymes and reagents employed is required to avoid interference with downstream enzymatic reactions. Innovations and process design can permit consolidation of the number of overall steps. For instance, the use of cap analogs in IVT can eliminate the need for a separate RNA capping reaction.^17^ However, a purification step is still required before the linearized template DNA can be used in IVT. In addition, for workflows that require enzymatic capping, the IVT product is usually purified before it can be used in RNA capping reactions.

Previously, we reported the use of immobilized SNAP-tagged poly(A) polymerase and immobilized SNAP-tagged T4 DNA ligase to Sera-Mag™ beads in direct RNA-seq, highlighting a one-pot workflow that generates a sequencing-ready library for direct loading onto a MinION flow cell. This streamlined immobilized workflow significantly lowered the input threshold for direct meta-transcriptomic RNA-seq and exhibited an improved transcriptome profiling.^22^ SNAP-tag is a small polypeptide (20 kDa) that can easily be engineered into recombinant proteins. Immobilization via the covalent SNAP-benzylguanine linkage involves gentle conditions that are compatible to preserving enzyme activity (**Supplementary** Fig.1). We also showed that manipulation of the substrate surface chemistry, such as the addition of a hydrophilic polymer (e.g. polyethylene glycol) can significantly impact enzyme properties.^22–24^

In this work, we show that solid-phase catalysis can also be used to streamline *in vitro* synthesis of capped mRNA and reduce sample loss by eliminating purification steps. For example, plasmids linearized by immobilized BspQI can be used directly in IVT reactions without purification. In addition, some of the key enzymes in mRNA synthesis workflows exhibit excellent activity when immobilized and are more resilient to heat than their soluble counterparts. We also demonstrate that some enzymes can be used repeatedly in their immobilized form. We further show that co-immobilizing an FCE::T7RNAP fusion with vaccinia cap 2′-O-methyltransferase facilitates efficient Cap-1 mRNA synthesis in a single-step, single-vessel workflow. All in all, we show that processes requiring multiple enzymatic reactions in series can benefit from enzyme immobilization (**Fig.1**), and that solid-phase enzymatic catalysis can help devise efficient, and scalable IVT processes.

**Figure 1.**
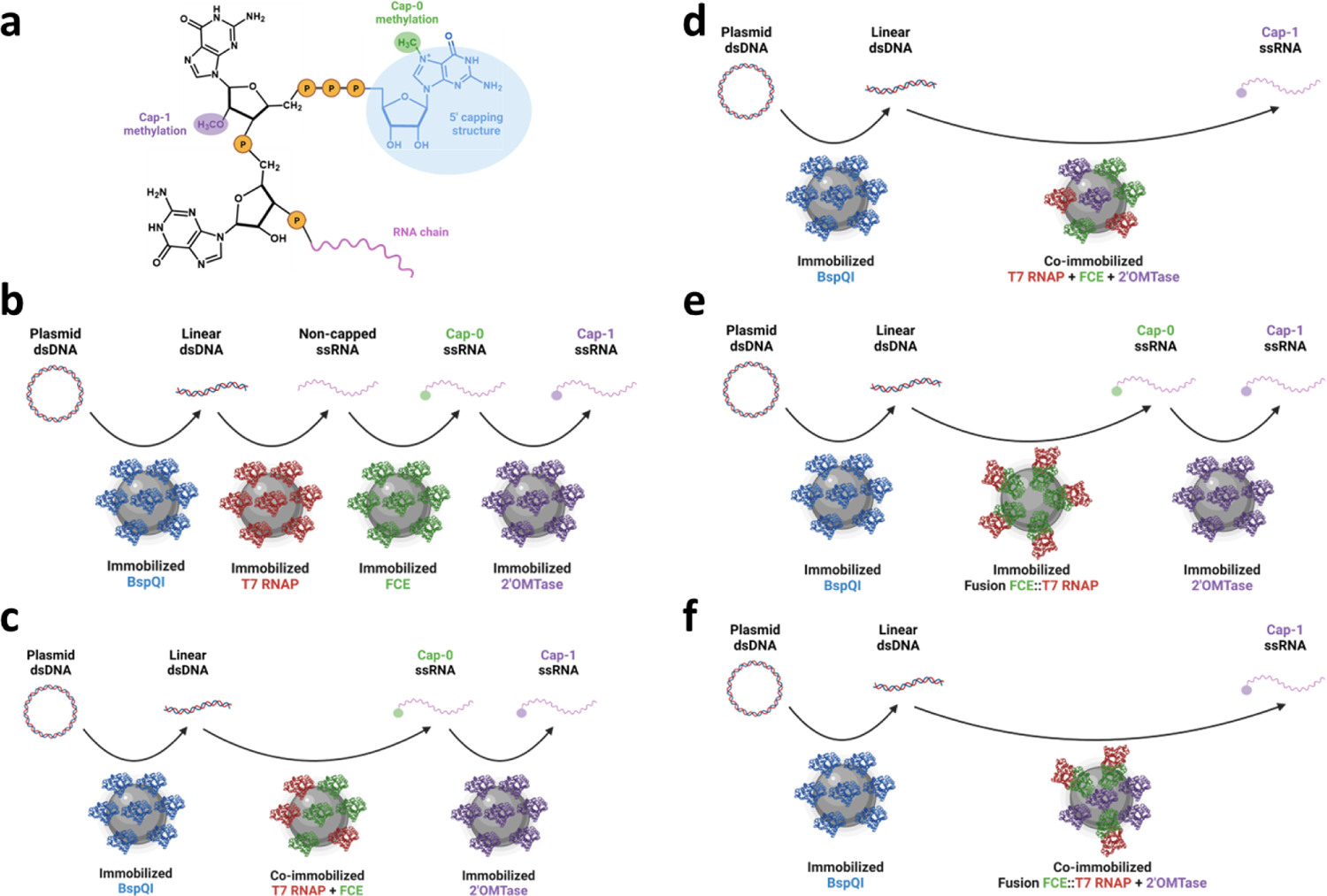
Schematic representations of capped RNA synthesis workflows using immobilized enzymes tested in this work. **(a)** The 5′ Cap-1 structure. The Cap-0 structure lacks the methyl group at the 2′-O position of the first nucleotide (in purple). **(b)** A continuous workflow for *in vitro* mRNA synthesis in consecutive reactions involving immobilized enzymes: BspQI generates linear DNA template, T7 RNAP catalyzes *in vitro* transcription, FCE yields Cap-0 product, and 2′OMTase achieves Cap-1 methylation. **(c)** A workflow that involves T7 RNAP and FCE co-immobilized onto the same microbeads to achieve single reaction mixture co-transcriptional capping. **(d)** A workflow that uses T7 RNAP, FCE and 2′OMTase co-immobilized onto the same microbeads to achieve one-pot co-transcriptional capping and Cap-1 methylation. **(e)** A workflow conducted by immobilizing an FCE::T7 RNAP fusion protein to improve coupled *in vitro* RNA synthesis and 5′ capping reactions. **(f)** A workflow that utilizes the FCE::T7 RNAP fusion protein and 2′OMTase co-immobilized onto the same microbeads to achieve one-pot co-transcriptional capping and Cap-1 methylation.

## RESULTS

### DNA template linearization using immobilized BspQI results in higher IVT output per input DNA

IVT requires DNA templates which are usually linearized plasmid DNA such that T7 RNA polymerase runs off the cleaved end of the template sequence to generate RNA transcripts that terminate at the desired position. Plasmid linearization is commonly done using a Type IIS restriction endonuclease (RE), such as BspQI, to create a “scarless” template that does not contain RE recognition sequence.^25^ The linearized DNA is then purified prior to the IVT reaction to avoid undesirable reactions such as off-target cleavage of the plasmid DNA by the restriction enzyme during storage. The purification step adds time, complexity and introduces sample loss. Thus, we aimed to create a new workflow where plasmid linearization can be incorporated in-line with IVT by using immobilized BspQI in an IVT-compatible buffer so that the reaction medium containing linearized DNA template can be directly used in IVT reactions after removing the immobilized BspQI.

We first created a fusion gene expressing BspQI, a 6xHis-tag and a SNAP-tag. The SNAP-tagged BspQI fusion protein was purified and immobilized onto benzylguanine-coated magnetic beads (BspQI@BG). The estimated protein load on beads was 20 mg g^-1^ carrier (**Supplementary** Fig.2, **Supplementary Table 1**). The activity of the immobilized enzyme was first evaluated using a FAM-labeled oligonucleotide (60-bp) containing one BspQI restriction site (**Fig.2a**, **Supplementary Table 2**). Complete cleavage of 700 nM of the DNA was achieved using 125 nM of either free or immobilized BspQI. At lower enzyme concentrations (15.62 and 7.81 nM), immobilized BspQI@BG showed approximately 15% lower activity compared to free BspQI (**Supplementary** Fig.3).

**Figure 2.**
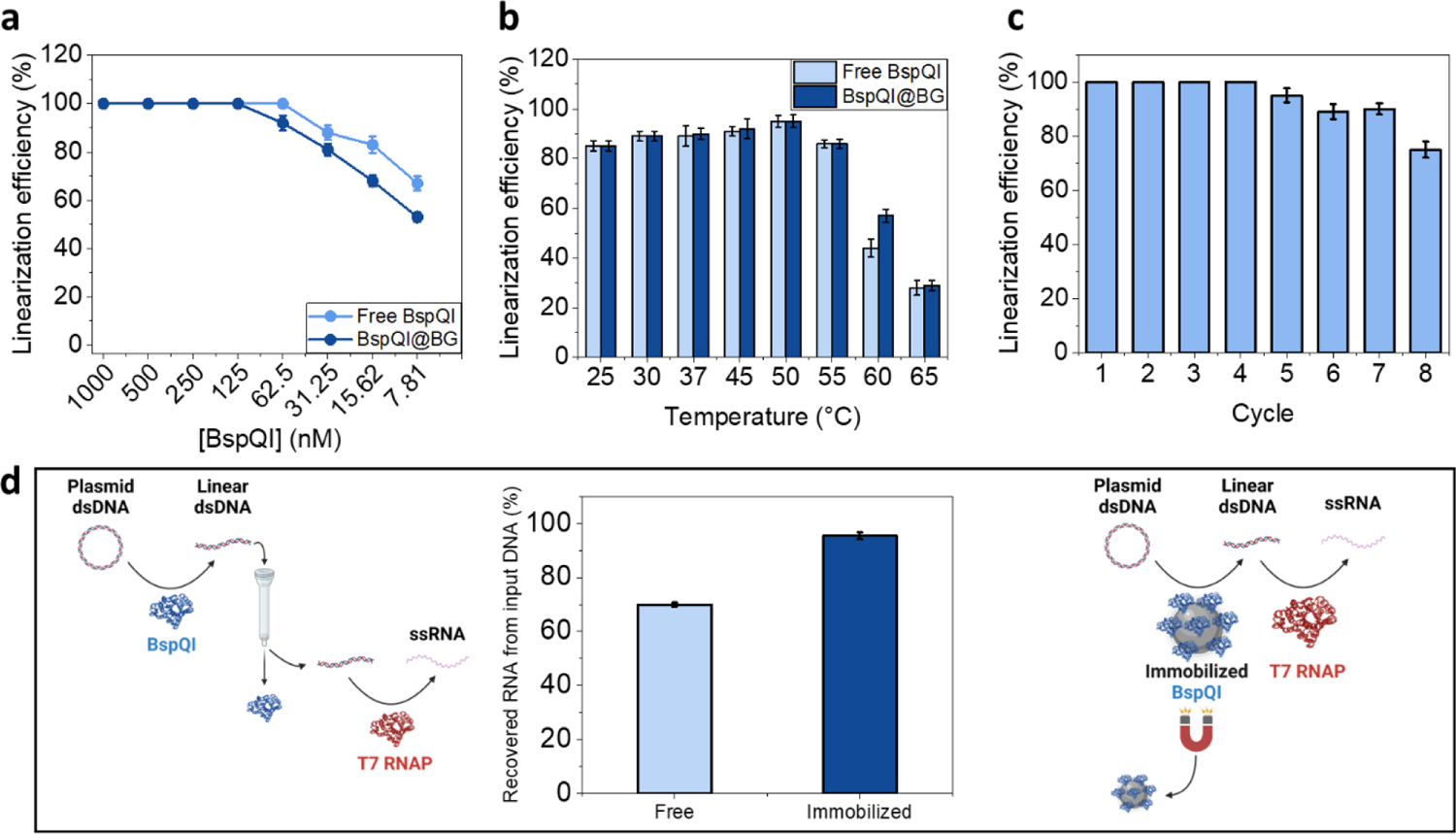
Evaluation of BspQI immobilization. **(a)** Immobilization efficiency. 700 nM of a 60-bp FAM-labeled oligonucleotide DNA was incubated with decreasing concentrations of free or immobilized BspQI at 50°C for 1 h. **(b)** Cleavage efficiency at different temperatures. Reactions were carried out at 25 to 65°C using 1 µM of free or immobilized BspQI (BspQI@BG). **(c)** Repeated use of immobilized BspQI. 1 µM of BspQI@BG was subjected to 8 cycles of 1 h, 50°C reaction on 700 nM of the of 60-mer FAM labeled oligonucleotide with thorough washing in RNA polymerase buffer between each cycle. **(d)** Immobilized BspQI facilitates higher RNA yield per input DNA. IVT of a 173 nt RNA was performed on a template based on pRNA21 was done using 0.15 µM of free or immobilized BspQI. The left panel depicts DNA linearization using free BspQI, followed by column purification and IVT. The right panel depicts DNA linearized using immobilized BspQI, which is subsequently removed with a magnet. Without purification, linearized DNA was used directly in IVT. The graph shows the RNA yield in terms of µg of transcribed RNA per µg of input circular DNA template. Average values of triplicate experiments are shown. Error bars represent one standard deviation.

Free BspQI remains highly active across a broad range of temperatures, supporting greater than 80% cleavage of DNA substrates at temperatures of 25 to 37°C, and ∼ 100% at 45 and 50°C (**Fig.2b**, **Supplementary** Fig.4), with a substantial activity drop at 60°C and above. Immobilized BspQI exhibited activity comparable to its free counterpart between 25 to 55°C while retaining higher activity at 60°C. Immobilized BspQI further exhibited substantial thermostability when it was re-used in cycles of 1 hour-reactions at 50°C (**Fig.2c**, **Supplementary** Fig.5), maintaining 100% activity in the first 4 cycles of reaction. Cleavage activity started to decrease at cycle 5 while exhibiting 70% activity in the 8^th^ cycle.

Next, we tested BspQI activity on a 60-bp DNA oligo in two reaction buffers: NEBuffer 3.1, as recommended by the manufacturer, and RNA polymerase buffer, with the goal of using the cleavage product directly without the need of purification or buffer exchange. We found that both free and immobilized BspQI are more active in RNA polymerase buffer than in NEBuffer 3.1 (**Supplementary** Fig.3), permitting direct interface of BspQI digestion with IVT.

To demonstrate the direct use of immobilized BspQI digestion products in IVT, a 173 nt RNA was transcribed using pRNA21 as template (**Supplementary Table 3**). The IVT reactions were performed using (1) a traditional workflow that involves free BspQI followed by column purification and a separate IVT reaction (**Fig.2d**, left panel), or (2) a workflow where the template DNA was linearized by immobilized BspQI and directly used in IVT (**Fig.2d**, right panel). We found that the DNA purification step resulted in about 30% loss of linearized DNA product in the traditional workflow (**Fig.2d**, left panel, **Supplementary** Fig.6). In contrast, DNA loss was not observed using immobilized BspQI (**Fig.2d**, right panel). We further assessed the impact of the workflow on IVT yield. We found that with the same amount of input DNA, the RNA yield was approximately two-fold higher using the immobilized BspQI workflows compared to the free enzyme workflow (**Fig.2d**, middle panel). Therefore, the use of immobilized BspQI can facilitate consolidation of plasmid linearization and IVT for a streamlined and efficient mRNA synthesis process.

### *In vitro* transcription using immobilized T7 RNA polymerase

To try to further streamline the mRNA synthesis process, we created a SNAP-tagged T7 RNA polymerase (T7 RNAP) construct and immobilized the purified SNAP-tagged T7 RNAP protein on BG-magnetic beads. We tested T7 RNAP immobilization on beads with polyethylene glycol (PEG) coating (T7@PEG) and without (T7@BG), as we previously demonstrated that some enzymes may require a more hydrophilic environment to retain their activity in solid-phase.^22, 26^ We achieved an estimated protein load on beads of ∼ 20 mg g^-1^ carrier (**Supplementary** Fig.2, **Supplementary Table 1**).

Using BspQI linearized-pRNA21 DNA as template, a time course of IVT showed that free T7 RNAP has a specific productivity (SP, **Supplementary Table 4**) about 3-fold higher than T7 RNAP immobilized on PEG_750_-coated beads (T7@PEG), which exhibits a slightly higher yield over T7 RNAP immobilized on uncoated beads (T7@BG) (**Fig.3a**). This suggests that T7 RNAP activity may be sensitive to the hydrophobic environment presented at the bead surface compared to a more hydrophilic bead surface, as previously reported for other enzymes.^22, 26^ T7 RNAP is most active at 37°C in free and immobilized forms (**Fig.3b**). While free T7 RNAP is highly active at 45°C (∼80% activity), T7@BG and T7@PEG retained only 10% activity. Immobilized T7 RNAP exhibits lower reusability when compared to immobilized BspQI, where T7@BG lost 20% or more of its activity in each cycle (**Fig.3c**). Attempts to improve the re-usability of immobilized T7 RNAP are underway.

**Figure 3.**
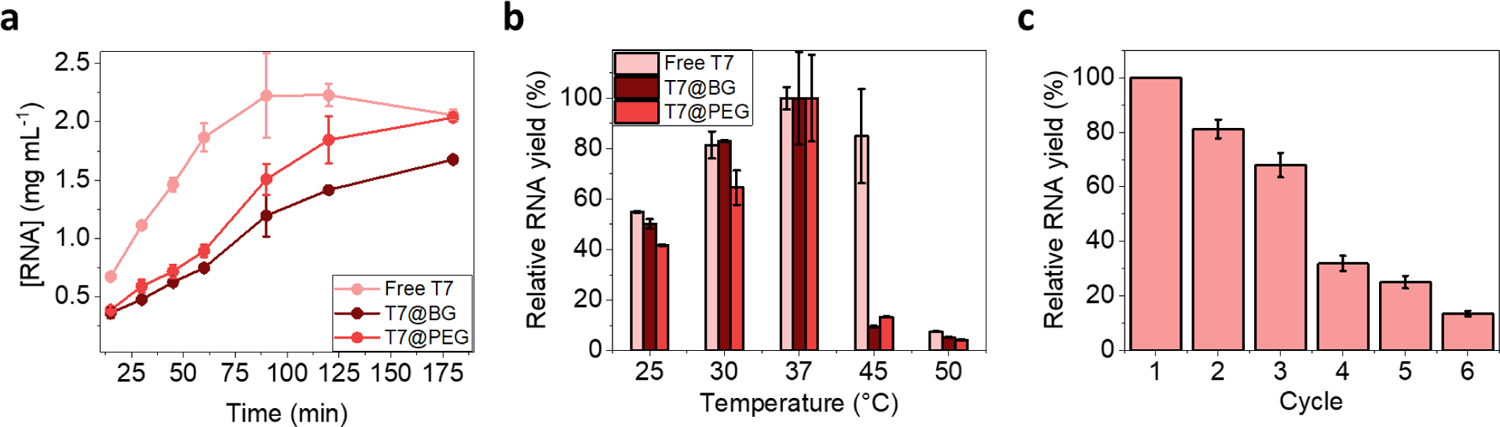
*In vitro* transcription using immobilized T7 RNA polymerase. **(a)** Time course of IVT using 0.17 µM of free T7 RNAP, T7@BG and T7@PEG at 37°C. **(b)** Relative RNA yield (%) compared to the maximum yield for each enzyme preparation (free T7, T7@BG and T7@PEG). IVT reactions were incubated at indicated temperatures (25 to 50°C) for 2 h. **(c)** Relative RNA yield (%) of T7@BG compared to the maximum yield achieved in cycle 1. The immobilized enzyme was reused in subsequent cycles of 2 h at 37°C after thorough washing in RNA polymerase buffer.

### mRNA Cap-0 and Cap-1 incorporation using immobilized Faustovirus RNA capping enzyme and vaccinia cap 2′-O-methyltransferase

Faustovirus capping enzyme (FCE) is a single-subunit RNA capping enzyme that exhibits high specific activity, broad temperature range, and possesses a three-domain fold similar to the larger subunit of VCE (**Fig.4a**).^27^ To incorporate RNA capping in an immobilized enzyme workflow, we produced SNAP-tagged versions of FCE and vaccinia RNA cap 2′-O-methyltransferase (2′OMTase) and immobilized them on BG- and PEG_750_-coated magnetic beads. Both SNAP-tagged enzymes were immobilized to the beads at high efficiency and exhibited higher enzyme activity on PEG_750_-coated beads (FCE@PEG and 2′OMTase@PEG) (**Supplementary** Fig.2, **Supplementary Table 1**).

Using a capillary electrophoresis based-assay,^28^ the RNA capping activity of free and immobilized FCE on a 5′ triphosphate, 3′ FAM-labeled RNA oligonucleotide (ppp25mer) was evaluated (**Supplementary Table 2**). **Fig.4b** (and **Supplementary** Fig.7) shows the activity of FCE at various temperatures. Consistent with previous results,^27^ FCE is highly active across a broad range of temperatures from 25 to 55°C. Immobilization on BG-PEG_750_-coated magnetic beads (FCE@PEG) improved FCE activity at the higher end of the temperature range (55-65°C). FCE@PEG exhibits >90% capping activity up to 4 cycles of use at 45°C (**Fig.4c**, **Supplementary** Fig.8). Therefore, this immobilization strategy is a promising approach for scaling FCE-mediated Cap-0 mRNA modification reactions.

The enzymatic activity of immobilized 2′OMTase was assayed using a Cap-0 RNA oligo and LC-MS/MS as readout. The immobilized 2′OMTase was highly active up to 4 cycles of use at 37°C or 5 cycles at 45°C (**Fig.4d**). The enzyme lost ∼20% each cycle after the 4^th^ cycle at 37°C and ∼10% after the 5^th^ cycle when used at 45°C.

Capping efficiency was further tested using a purified 1653 nt transcript at 45°C (**Supplementary Table 3**). Both free and immobilized FCE achieved near to 100% Cap-0 yield of the full mRNA transcript (**Fig.4e**). Cap-1 modification of this transcript neared 100% when using free and immobilized 2′OMTase were added under these same conditions. Taken together, immobilized FCE and vaccinia cap 2′-O-methyltransferase facilitate effective RNA 5′ cap modification. The compatibility to biocatalyst recycling and wider range of reaction temperature may allow for further innovation in mRNA manufacturing processes.

### Co-transcriptional Cap-0 and Cap-1 RNA synthesis using immobilized enzymes

To evaluate if the immobilized enzymes can achieve capped RNA synthesis in a one-pot format, individually immobilized enzymes (each bead has only one of the enzymes immobilized) and co-immobilized enzymes (each bead has a combination of different enzymes immobilized were tested in co-transcriptional conditions. Transcription yield and cap incorporation was evaluated for the 174 nt pRNA21 transcript. Co-transcriptional capping reactions were performed at 45°C to take advantage of the higher FCE activity (Fig.4).

**Figure 4.**
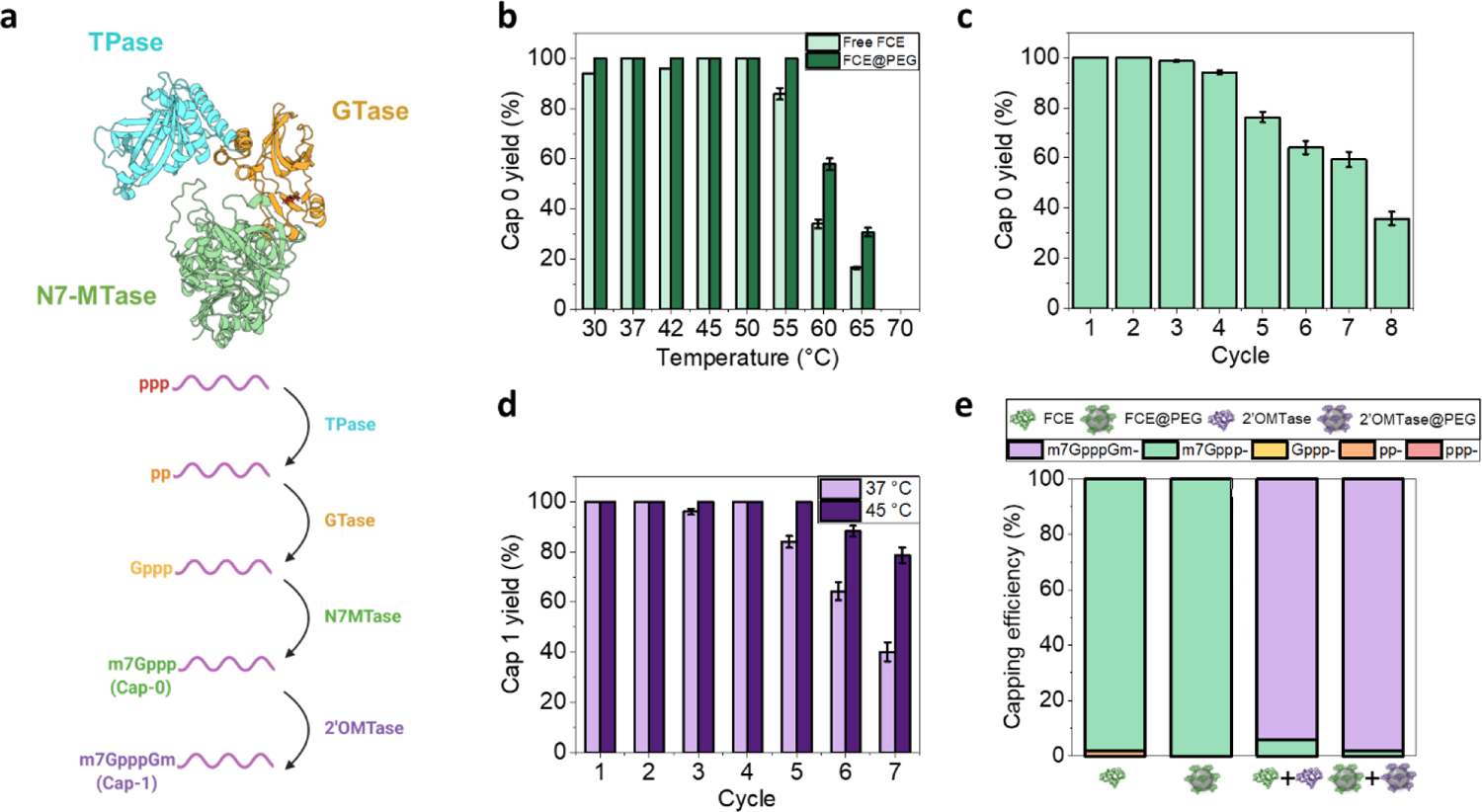
*In vitro* Cap-0 and Cap-1 mRNA capping using immobilized FCE and 2′OMTase. **(a)** Top, an AlphaFold2 prediction of the FCE structure. The functional domains are colored: cyan, triphosphatase (TPase) domain; gold, guanylyl transferase (GTase) domain; green, N7-methyl transferase (N7-MTase) domain. Bottom, a schematic of the sequential enzymatic reactions to form Cap-0 (m7Gppp-) and Cap-1 (m7GpppGm-). ppp-: 5′ triphosphate RNA. pp-: 5′ diphosphate RNA. Gppp-: 5′ guanylyl-triphosphate RNA. m7Gppp-: Cap-0 RNA. m7GpppGm-: Cap-1 RNA. **(b)** RNA capping activity of free and immobilized FCE (FCE@PEG). 0.5 µM of ppp25mer RNA oligo was incubated with 0.2 µM of free FCE or FCE@PEG at designated temperatures (from 30 to 70°C) for 30 min. The reactions were quenched and analyzed by capillary electrophoresis. **(c)** Repeated reuse of FCE@PEG. 0.5 µM of the ppp25mer RNA was incubated with 0.2 µM of FCE@PEG in 8 reaction cycles of 30 min at 45°C. The FCE@PEG beads were washed thoroughly in FCE capping buffer between cycles. Cap-0 yield was represented as percentage relative to maximum yield achieved in cycle 1. **(d)** Repeated reuse of 2′OMTase@PEG. 0.5 µM of Cap-0 25mer RNA was incubated with 0.2 µM of 2′OMTase@PEG in seven 30 min reaction cycles at 37°C or 45°C. The 2′OMTase@PEG beads were washed thoroughly in FCE capping buffer between cycles. Cap-1 yield was represented as percentage relative to maximum yield achieved in cycle 1. **(e)** Performance of immobilized FCE and 2′OMTase on long RNA. 1 µM of a 1.7 kb FLuc *in vitro* transcript was incubated with 0.2 µM of free FCE, FCE@PEG or a combination of free FCE plus free 2′OMTase or FCE@PEG plus 2′OMTase@PEG. Enzymes in free or immobilized forms are represented as stylized structural models (see figure legends).

To evaluate the performance of immobilized enzymes in generating Cap-0 RNA in a one-pot format, 0.5 µM of T7 RNAP and 0.5 µM of FCE were incubated with 0.5 µM of BspQ1-linearized pRNA21. We found that individually immobilized T7@BG and FCE@BG returned low capping efficiency (5% Cap-0) with a transcription yield similar to when free T7 RNAP and FCE were used (**Fig.5a**, Rxn 1). T7 RNAP and FCE immobilized on PEG_750_-coated beads performed better at ∼30% Cap-0 (Rxn 2). Co-immobilizing T7 RNAP and FCE (T7RNAP+FCE@BG) further improved the extent of Cap-0 incorporation to ∼50% with a similar yield of transcription (Rxn 3). However, co-immobilizing the two enzymes on PEG_750_-coated beads (T7 RNAP+FCE@PEG) decreased Cap-0 incorporation to ∼30% (Rxn 4).

**Figure 5.**
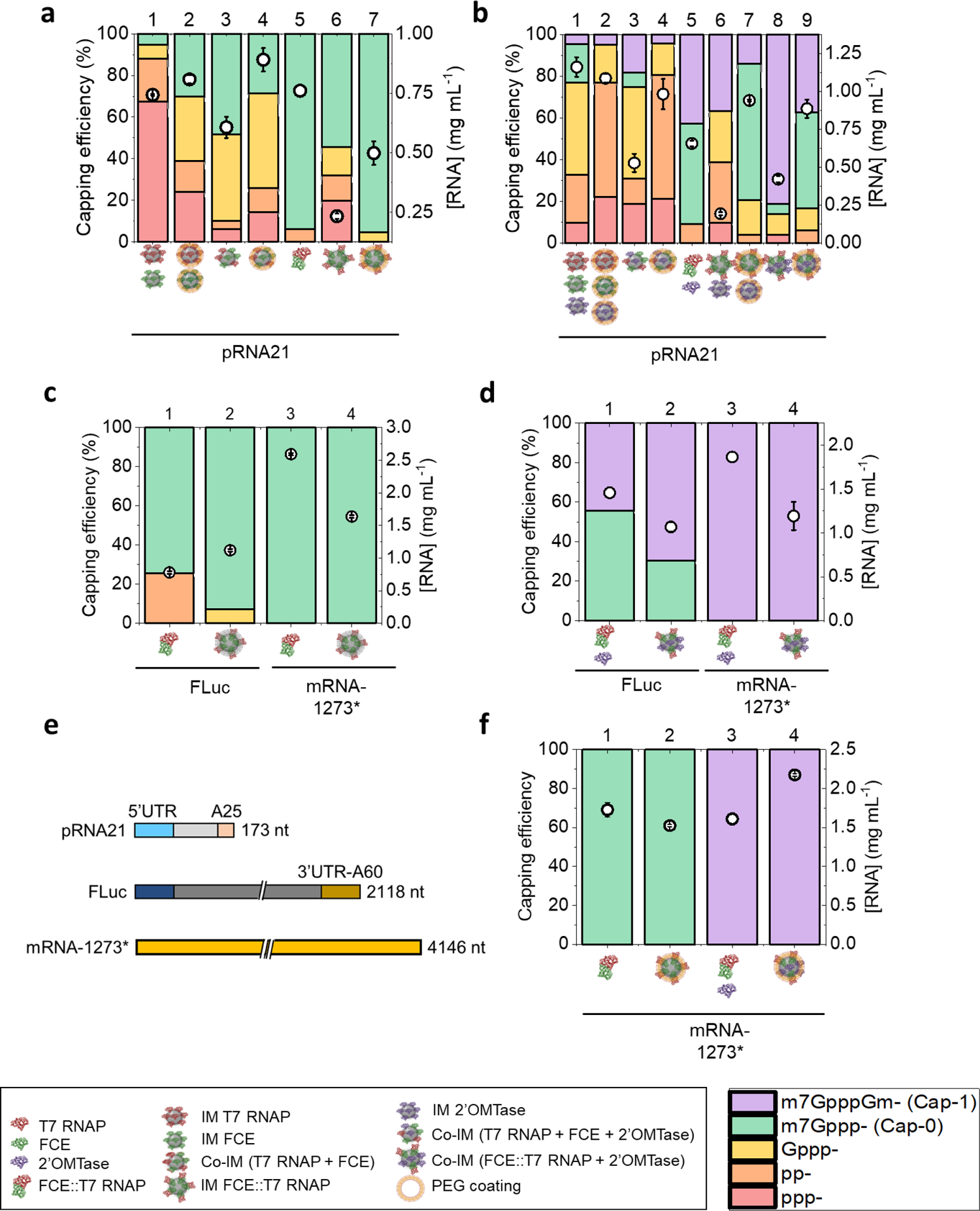
Single reaction mixture capped RNA synthesis using free and immobilized enzymes (0.5 µM). Transcription yield (white circles) in mg mL^-1^, is indicated on the right-Y axes. In all cases, reactions were carried out at 37°C for 90 minutes, followed by an incubation at 45°C for 30 minutes. Cap-0 **(a)** and Cap-1 **(b)** incorporation (%, left-Y axes, stacked columns) of a 173 nt transcript from a PCR amplicon derived from pRNA21. Enzymes used (free or immobilized) were indicated as stylized structural depictions (see keys in the middle panel). Cap-0 **(c)** and Cap-1 **(d)** incorporation (%, left-Y axes, stacked columns) of the 1.7 kb FLuc transcript, and the 4.1 kb transcript derived from the mRNA-1273* sequence using free or immobilized enzymes. **(e)** Schematic representation of the three mRNA transcripts obtained in this experimental section. **(f)** Cap-0 and Cap-1 incorporations using free or immobilized enzymes and substituting UTP with m_1_ψTP in the IVT reaction.

To try to further improve Cap-0 incorporation using the immobilized enzyme platform, we generated a SNAP-tagged version of an FCE::T7RNAP fusion protein (manuscript submitted concurrently) and immobilized it to BG-magnetic beads. In nature, RNA capping enzymes invariably associate with the mRNA-transcribing RNA polymerase ^29^ such that nascent mRNAs are capped co-transcriptionally when ∼20 nt have been synthesized.^30–34^

Similar to the non-SNAP-tagged version (manuscript submitted concurrently), the SNAP-tagged FCE::T7RNAP fusion was highly efficient in generating >90% Cap-0 *in vitro* transcript as free enzyme (**Fig.5a**, Rxn 5). The use of PEG_750_-coated beads increased Cap-0 incorporation to ∼95% with a yield of ∼0.5 mg mL^-1^ (Rxn 7). Interestingly, when immobilized on BG-beads, however, the fusion protein only generated 50 % Cap-0 transcript with a low transcription yield (∼0.25 mg mL^-1^) (Rxn 6).

We next worked toward enzymatic co-transcriptional mRNA capping that includes Cap-1 modification. To that end, we investigated immobilized Cap 2′OMTase could support enzymatic co-transcriptional mRNA capping in an one-pot format. **Fig.5b** shows that using individually immobilized T7 RNAP, FCE and 2′OMTase with or without PEG_750_ coating produced similar transcription yield but decreased the extent of Cap-1 to ∼5% (Rxn 1 and 2). Co-immobilizing the three enzymes on BG-beads with or without PEG_750_ coating only generated ∼20% or ∼5%, respectively, Cap-1 transcripts (Rxn 3 and 4).

Next, we combined free FCE::T7RNAP fusion with 2′OMTase in a single reaction mixture and observed ∼40% Cap-1 transcript (Rxn 5). Cap-1 incorporation was maintained at ∼40% using individually immobilized FCE::T7RNAP and 2′OMTase (FCE::T7RNAP@BG and 2′OMTase@BG) but transcript yield decreased to 0.2 mg mL^-1^ (Rxn 6). Using PEG_750_-coated beads improved transcript yield but did not improve Cap-1 incorporation (Rxn 7). Co-immobilizing the FCE::T7RNAP fusion and 2′OMTase (FCE::T7RNAP+2′OMTase@BG) greatly improved Cap-1 incorporation to ∼80% with 0.4 mg mL^-1^ transcript yield (Rxn 8). Co-immobilizing on PEG_750_-coated beads (FCE::T7RNAP+2′OMTase@PEG, Fig.5b, Rxn 9) decreased the level of Cap-1 incorporation to ∼40%, even though individually immobilized FCE::T7RNAP (**Fig.5a**, Rxn 7) and 2′OMTase (**Fig.4e**) performed better on PEG_750_-coated beads.

We next evaluated the performance of the most promising immobilized biocatalysts FCE::T7@PEG, and FCE::T7RNAP+2′OMTase@BG for synthesizing Cap-0 or Cap-1 RNA in an one-pot format for different DNA templates (**Fig.5e**, **Supplementary Table 3, Supplementary** Fig.9). As shown in **Fig.5c**, FCE::T7@PEG generated ∼95% of FLuc *in vitro* transcripts containing the Cap-0 structure (Rxn 2), on par with the performance of its free counterpart (Rxn 1). FCE::T7@PEG generated 100% Cap-0 modified 4.1 kb *in vitro* transcript based on mRNA-1273 of the original Moderna SARS-CoV-2 mRNA vaccine (mRNA-1273*) with a transcript yield of ∼1.6 mg mL^-1^ (Rxn 4). This is similar to the level of Cap-0 incorporation by free FCE::T7RNAP albeit a lower yield (Rxn 3).

For single reaction mixture enzymatic Cap-1 mRNA synthesis (**Fig.5d**), FCE::T7RNAP and 2′OMTase co-immobilized on uncoated BG-beads (FCE::T7RNAP+2′OMTase@BG) generated more Cap-1 FLuc RNA (70%, Rxn 2) compared to free FCE::T7RNAP and 2′OMTase (45%, Rxn 1). The co-immobilized enzymes FCE::T7RNAP+2′OMTase@BG generated 100% Cap-1 mRNA-1273* at ∼1.6 mg mL^-1^ transcript yield (Rxn 4), respectively, compared to 100% Cap-1 at ∼2.5 mg mL^-1^ to its free counterpart (Rxn 3).

Finally, we demonstrate that the substitution of uridine by non-canonical m_1_-pseudo-uridine (m_1_-ψ) generates ∼1.7 and ∼1.5 mg mL^-1^ Cap-0 mRNA-1273* when using free FCE::T7 RNAP and FCE::T7@PEG (**Fig.5f**, Rxn 1 and 2), respectively. Cap-1 mRNA-1273* was also obtained with free FCE::T7RNAP and 2′OMTase and its BG-immobilized counterpart at ∼1.6 and ∼2.2 mg mL^-1^ (Rxn 3 and 4), respectively, using m1-ψ in the reaction.

## DISCUSSION

mRNA synthesis is a multi-stage process that involves DNA template preparation, *in vitro* transcription, 5′ cap and 3′ poly(A) tail incorporation. Currently, RNA capping can be performed using RNA capping enzymes post-transcriptionally or cap analogs co-transcriptionally. Although the use of cap analogs allows for one-pot capped RNA synthesis, only a small fraction of input cap analog is incorporated into the RNA molecule, leaving most of the cap analog unusable.^17, 35–37^

Current workflows require separate enzymatic reactions for plasmid template linearization, IVT using templated poly(A) tail incorporation, followed by enzymatic RNA capping with intermediate purifications, dilutions, or buffer exchanges.^38, 39^ Streaming the manufacturing can help reduce cost and make mRNA manufacturing more environmentally sustainable. To that end, we engineered major mRNA synthesis enzymes to be immobilized to solid supports and assessed the immobilized enzymes′ performance in terms of enzyme activity, reusability, and in sequential and combined workflow settings.

We report that the immobilized enzymes exhibit similar or improved enzymatic properties compared to their solution counterparts, and that the use of immobilized enzymes in mRNA synthesis offers several advantages over the current multi-stage workflows. For example, immobilized BspQI facilitates direct interface of template linearization to IVT, eliminating the linearized plasmid purification step and improving the RNA yield per input circular plasmid. Eliminating the purification step can also prevent damage to the plasmid DNA (such as nicking) or carryover of chaotropic and organic agents used in linearized DNA purification into IVT reactions. FCE and vaccinia cap 2′-O-methyltransferase exhibit excellent enzyme activity upon repeated use in reaction cycles, potentially making mRNA synthesis a more sustainable process.

In eukaryotic mRNA biogenesis, mRNA capping is directly coupled to RNA synthesis through interactions of RNA capping enzyme with the C-terminal domain repeat sequences of RNA polymerase II.^31, 33^ Additional evidence supports capping enzyme-RNAP interactions separate from the CTD, and the positioning of the TPase and GTase active sites of the RNA capping enzyme at the vicinity of the RNA exit tunnel of the RNA polymerase.^29^ Capping of nascent transcripts before their release from RNAP is an efficient natural mechanism to install the cap structures. To reproduce co-transcriptional enzymatic capping in vitro, we showed that high Cap-0 and Cap-1 mRNA (up to 95%) can be achieved using an FCE::T7RNAP fusion protein (manuscript produced concurrently). We further show in this report that co-immobilized FCE::T7RNAP fusion-cap 2′OMTase is highly effective in generating Cap-1 RNA in a one-pot co-transcriptional capping format.

In addition, immobilized enzymes could exhibit superior properties such as higher thermostability/thermoactivity, reusability and the capability of adapting a continuous flow format, as already demonstrated for many relevant enzyme-catalyzed reactions in chemical and pharmaceutical industries.^40–44^ These attributes can be advantageous for scale up and scale out approaches to increasing the capacity and specialization of mRNA manufacturing process. More research on immobilization substrates, surface chemistry and reaction format (such as continuous flow) can further improve the performance and the applicability of immobilized enzymes in processes such as mRNA manufacturing and molecular diagnostic applications.

In conclusion, we show that solid-phase catalysts can streamline and open new possibilities for mRNA manufacturing. Elimination of intermediate purification steps and conducting one-pot multiple enzymatic reactions through enzyme immobilization and the use of novel enzyme fusions may help increase the environmental sustainability of mRNA synthesis at scale by reducing waste and reliance on cap analogs manufactured by organic synthesis.

## METHODS

### Materials

Carboxylate-Modified Magnetic Beads (Sera-Mag^TM^; 50 mg mL^-1^) and chromatography columns HiTrap® diethylaminoethyl (DEAE), HisTrap® Ni Sepharose and HiTrap® Heparin were purchased from Cytiva (Marlborough, MA, USA). Magnetic separation racks, Benzylguanine-NH2 (BG-NH_2_), dithiotreitol (DTT), all reaction buffers (10x concentration), yeast inorganic pyrophosphatase (100 U mL^-1^), murine RNase inhibitor (40,000 U mL^-1^), ribonucleotide solution mixture (rNTP mix: 25 mM rATP, 25 mM rGTP, 25 mM rCTP, 25 mM rUTP), S-adenosylmethionine (SAM, 32 mM), guanosine triphosphate (GTP, 10 mM), Monarch^®^ PCR & DNA Cleanup Kit, Monarch^®^ RNA Cleanup Kit were from New England Biolabs (Ipswich, MA, USA). 2-(N-morpholino)ethanesulfonic acid (MES) buffer, N-hydroxysuccinimide (NHS), diethanolamine, tris(2-carboxyethyl)phosphine (TCEP), Turbo DNase I (2 U µL^-1^), ethylenediaminetetraacetic acid solution (EDTA, 0.5 M), Nanodrop^TM^ 8000 were purchased from ThermoFisher Scientific (Waltham, MA, USA). Methoxypolyethylene glycol amine (PEG_750_), Tris-HCl, BisTris, NaCl, imidazole, L-Arginine were purchased from Sigma-Aldrich (Burlington, MA, USA), sodium dodecylsulfate solution (SDS, 20%) was purchased from Teknova (Mansfield, MA, USA).

### BspQI activity assay

Free and BG-coated beads immobilized BspQI (BspQI@BG) activities were assayed at different enzyme concentrations and reaction temperatures towards a chemically synthesized double-stranded DNA substrate (60-mer). A 1:9 mixture of FAM-labeled 60-mer (contains a FAM-labeled at the 5′ end of the top strand) and unlabeled 60-mer (0.7 µM) was incubated with serial dilutions of free BspQI or BspQI@BG at 50°C for 1 hour in NEBuffer r3.1 (10 mM NaCl, 5 mM Tris-HCl (pH 7.9), 1 mM MgCl_2_, 10 µg mL^-1^ recombinant albumin) or RNA polymerase buffer (4 mM Tris-HCl (pH 7.9), 0.6 mM MgCl_2_, 0.1 mM DTT and 0.2 mM spermidine) (**Fig.2a**, **Supplementary** Fig.2). After incubation, reactions were stopped by 10 mM EDTA, and analyzed by capillary electrophoresis using an Applied Biosystems 3730xl DNA Analyzer (ThermoFisher).^45^ To investigate the effect of reaction temperature on BspQI@BG, the assay was repeated at different reaction temperatures (25, 30, 37, 45, 50, 55, 60 and 65°C) (**Fig.2b**, **Supplementary** Fig.3). Recycling assay of BspQI@BG was performed by pelleting magnetic beads with a magnetic separation rack, washing beads three times using 1x reaction buffer, and adding fresh substrate mixture. Recycling process was repeated for 8 times (**Fig.2c**, **Supplementary** Fig.4). BspQI activity was also tested on pRNA21 DNA template plasmid. 0.5 mg mL^-1^ pRNA21 was incubated with 0.15 µM BspQI or BspQI@BG at 50°C for 1 hour in NEBuffer r3.1. Linearization efficiency was qualitatively determined using 1.2% agarose gel electrophoresis and DNA concentration was quantitatively determined using Nanodrop^TM^ 8000 spectrophotometer (ThermoFisher). To investigate if DNA linearized by BspQI@BG can be used directly in IVT without purification, pRNA21 linearized using BspQI@BG in RNA polymerase buffer was directly used in IVT after separating from the magnetic beads using a magnetic separation rack. pRNA21 linearized using free BspQI in NEBuffer r3.1 was purified using Monarch® PCR & DNA Cleanup Kit before using in IVT.

### T7 RNA polymerase activity assay

In vitro transcription activity of SNAP-tagged T7 RNA polymerase in solution or immobilized on BG-coated (T7@BG), or BG-PEG_750_-coated magnetic beads (T7@PEG) were assayed and compared at different reaction times and temperatures. 0.03 mg mL^-1^ of pRNA21 linearized by soluble BspQI was added to IVT reactions containing 5 mM of each rNTPs, 2.5 10^-3^ U μL^-1^ yeast inorganic pyrophosphatase, 1 U μL^-1^ murine RNase inhibitor, and 0.17 µM free T7 RNAP, T7 RNAP@BG or T7 RNAP@PEG in RNA polymerase buffer (New England Biolabs). For reaction time courses, reactions were incubated at 37°C for 15, 30, 45, 60, 90, 120 and 180 minutes (**Fig.3a**). To investigate the reaction temperature, reactions were incubated at 25, 30, 37, 45 and 50°C for 1 hour (**Fig.3b**). Recycling assay was performed by pelleting the T7@BG magnetic beads with a magnetic separation rack, washing beads three times in 1x RNA polymerase buffer, and adding reaction premix containing fresh NTPs, yeast inorganic pyrophosphatase, murine RNase inhibitor and template DNA. Recycling process was repeated for 6 times (**Fig.3c**). After each reaction, the reaction mixture was incubated with Turbo DNase I (2 U) at 37°C for 30 minutes. The in vitro transcript was purified using Monarch^®^ RNA Cleanup Kit. RNA concentration was quantitatively determined using a Nanodrop^TM^ 8000 spectrophotometer.

### RNA capping and Cap-1 methylation activity assay

SNAP-tagged Faustovirus RNA capping enzyme and vaccinia cap 2′-O-methyltransferase were immobilized on BG-coated (FCE@BG and 2′OMTase@BG) and BG-PEG_750_-coated magnetic beads (FCE@PEG and 2′OMTase@PEG). RNA capping activity of the immobilized and solution forms of SNAP-tagged FCE were assayed using capillary electrophoresis.^46^ Briefly, 0.2 µM the immobilized or solution form of SNAP-tagged FCE was incubated with 0.5 µM of a 5′ triphosphate 3′ FAM-labeled RNA oligonucleotide (ppp25-mer; Bio-synthesis Inc.) in reactions containing 50 mM Tris-HCl (pH 8.0), 5 mM KCl, 1 mM MgCl_2_, 1 mM DTT, 0.1 mM SAM and 0.5 mM GTP at 30, 37, 42, 45, 50, 55, 60, 65 and 70°C for 30 minutes. Reactions were then quenched using 10 mM EDTA and 1% SDS and analyzed by capillary electrophoresis using an Applied Biosystems 3730xl DNA Analyzer (ThermoFisher) (**Fig.4b** and **Supplementary** Fig.7). Recycling assay of FCE@PEG was performed by pelleting magnetic beads with a magnetic separation rack, washing beads three times, and adding fresh substrate mixture. Recycling process was repeated for 8 times (**Fig.4c** and **Supplementary** Fig.8). Cap-1 methylation activity of the solution and immobilized forms of vaccinia cap 2′OMTase was assayed by incubating 0.2 µM of solution of immobilized form of SNAP-tagged 2′OMTase with 0.5 µM of Cap-0 25mer (m7Gppp25mer) in reactions containing 50 mM Tris-HCl (pH 8.0), 5 mM KCl, 1 mM MgCl_2_, 1 mM DTT and 0.1 mM SAM. Reactions were incubated for 30 minutes at 37 or 45°C. Enzymes were recycled by pelleting the 2′OMTase@PEG magnetic beads with a magnetic separation rack, washing three times with reaction buffer and adding fresh substrate and SAM. Recycling process was repeated for 7 times. Reactions were quenched using 10 mM EDTA and 1% SDS. The level of methylation was analyzed using intact LC-MS analysis (**Fig.4d**).^46^ To test if immobilized FCE and 2′OMTase are compatible to one-pot Cap-1 incorporation, 0.2 µM of free FCE, FCE@PEG or a combination of free FCE plus free 2′OMTase or FCE@PEG plus 2′OMTase@PEG was incubated with 1 μM of a firefly luciferase in vitro transcript (FLuc; 1.7 kb) under the same reaction conditions. The RNA was then purified using Monarch® RNA Cleanup Kit and the level of cap incorporation was evaluated using a RNase H-based intact LC-MS analysis (**Fig.4e**).^28^

### Co-transcriptional mRNA capping activity assay

16.5 nM of linearized template plasmid was added to IVT reactions in RNA polymerase buffer (New England Biolabs) containing 5 mM of each rNTPs, 0.4 mM SAM, 2.5 10^-3^ U μL^-1^ yeast inorganic pyrophosphatase, 1 U μL^-1^ murine RNase inhibitor, and 0.5 µM of indicated enzymes. Reactions were incubated at 37°C for 90 minutes and at 45°C for 30 minutes. The reactions were then incubated with Turbo DNase I (0.1 U μ^-1^) at 37°C for 30 minutes. The *in vitro* transcripts were purified using Monarch® RNA Cleanup Kit. RNA concentration was determined using a Nanodrop^TM^ 8000 spectrophotometer (ThermoFisher). The level of cap incorporation was evaluated using a hRNase 4-based intact LC-MS analysis.^46^ One-pot co-transcriptional capping reactions were carried out using different combinations of biocatalysts: (1) free SNAP-tagged T7 RNAP and FCE (T7+FCE); (2) free SNAP-tagged FCE::T7RNAP fusion protein (FCE::T7); (3) SNAP-tagged T7 RNAP and FCE immobilized on BG-coated magnetic (T7@BG+FCE@BG); (4) SNAP-tagged T7 RNAP and FCE co-immobilized on BG-coated magnetic beads ([T7+FCE]@BG); (5) SNAP-tagged FCE::T7RNAP fusion protein (FCE::T7@BG); (6) BG-PEG_750_-coated beads immobilized T7 RNAP and FCE (in different bead units) in one pot (T7@PEG+FCE@PEG); (7) SNAP-tagged T7 RNAP and FCE co-immobilized on BG-PEG_750_-coated beads ([T7+FCE]@PEG); (8) SNAP-tagged FCE::T7RNAP fusion protein immobilized on BG-PEG_750_-coated beads (FCE::T7@PEG); (9) free T7 RNAP, free FCE and free 2′OMTase (T7+FCE+2′OMTase); (10) free FCE::T7RNAP fusion protein and 2′OMTase (FCE::T7+2′OMTase); (11) SNAP-tagged T7 RNAP, FCE and 2′OMTase immobilized on BG-coated magnetic beads (T7@BG+FCE@BG+2′OMTase@BG); (12) SNAP-tagged T7 RNAP, FCE and 2′OMTase co-immobilized on BG-coated magnetic beads ([T7+FCE+2′OMTase]@BG); (13) SNAP-tagged FCE::T7RNAP and 2′OMTase immobilized on BG-coated magnetic beads (FCE::T7@BG+2′OMTase@BG); (14) SNAP-tagged FCE::T7RNAP and 2′OMTase co-immobilized on BG-coated magnetic beads ([FCE::T7+2′OMTase]@BG); (15) SNAP-tagged T7 RNAP, FCE and 2′OMTase immobilized on BG-PEG_750_-coated magnetic beads (T7@PEG+FCE@PEG+2′OMTase@PEG); (16) SNAP-tagged T7 RNAP, FCE and 2′OMTase co-immobilized on BG-PEG_750_-coated magnetic beads [T7+FCE+2′OMTase]@PEG); (17) SNAP-tagged FCE::T7RNAP and 2′OMTase immoilzed on BG-PEG_750_-coated magnetic beads (FCE::T7@PEG+2′OMTase@PEG); (18) SNAP-tagged FCE::T7RNAP and 2′OMTase co-immobilized on BG-PEG_750_-coated magnetic beads ([FCE::T7+2′OMTase]@PEG) (**Fig.5a,b**). Template DNA includes (1) pRNA21; (2) a synthetic FLuc construct; (3) mRNA-1273* (a 4.1 kb transcript based on the Moderna COVID19 mRNA vaccine sequence) (**Fig.5c,d**).^47^ mRNA-1273* DNA co-transcriptional capping was also carried out using m1-pseudo-uridine triphosphate (**Fig.4f**).

