## Supplemental Materials for "Streamlined DNA template preparation and co-transcriptional 5′ capped RNA synthesis enabled by solid-phase catalysis"

### Supplementary information

**TABLE OF CONTENTS**


**SUPPLEMENTARY METHODS**

Functionalization of Sera-Mag™ Beads with Benzylguanine (BG) and Polyethylene glycol 750 (PEG) ........ S3

Protein expression and purification ........ S3

SNAP-tagged proteins immobilization in BG Magnetic Beads ........ S4

Calculation of immobilization parameters ........ S4

**SUPPLEMENTARY FIGURES**

Supplementary Figure 1 ........ S5

Supplementary Figure 2 ........ S6

Supplementary Figure 3 ........ S8

Supplementary Figure 4 ........ S9

Supplementary Figure 5 ........ S10

Supplementary Figure 6 ........ S11

Supplementary Figure 7 ........ S12

Supplementary Figure 8 ........ S13

Supplementary Figure 9 ........ S14

**SUPPLEMENTARY TABLES**

Supplementary Table 1 ........ S15

Supplementary Table 2 ........................................................................................................ S16

Supplementary Table 3 ........................................................................................................ S17

Supplementary Table 4 ........................................................................................................ S20

Supplementary Table 5 ........................................................................................................ S21

#### Supplementary Methods

##### Functionalization of Sera Magnetic Beads with Benzylguanine (BG) and Polyethylene glycol 750 (PEG)

100 mg of Sera-Mag™ 50 mg mL$^{-1}$ were diluted in isopropanol/water 1:1 mixture (IPA/water 1:1) to a final concentration of 10 mg mL$^{-1}$. Supernatant was removed by using magnetic separation rack. Beads were washed three times with 50 mM MES buffer pH 6. Supernatant was removed, beads were resuspended in 50 mM MES buffer pH 6, 50 mM NHS to a final concentration of 10 mg mL$^{-1}$ and incubated for 3 hours at room temperature with rotation. In parallel, 10 mg of BG-$NH_2$ were dissolved in DMSO to a final concentration of 6.7 mg mL$^{-1}$. BG-$NH_2$ was diluted in IPA/water 1:1 to a final concentration of 1 mg mL$^{-1}$ (3.7 mM). The beads' supernatant was removed, and beads were washed three times with ice-cold IPA/water 1:1. The supernatant was removed, beads were resuspended in 3.7 mM BG-$NH_2$ solution and incubated for 4 hours at room temperature. The supernatant was removed. For non-PEG coated bead preparation, the beads were resuspended in 0.2 M diethanolamine in IPA/water 1:1 to a final concentration of 10 mg mL$^{-1}$ and incubated for 2 hours at room temperature. For PEG-coated beads preparation, instead, the beads were resuspended in 20 mM $PEG_{750}$ in IPA/water 1:1 to a final concentration of 10 mg mL$^{-1}$ and incubated for 24 hours at room temperature. The supernatant was removed, and the beads were resuspended in 0.2 M diethanolamine in IPA/water 1:1 to a final concentration of 10 mg mL$^{-1}$ and incubated for 2 hours at room temperature. For both non-PEG and PEG-coated beads, the supernatant was removed, and the beads were washed three times with water, and three times with IPA/water 1:1. The beads were resuspended in IPA/water 1:1 to a final concentration of 10 mg mL$^{-1}$.

##### Protein expression and purification

The plasmids encoding His- SNAP-tagged variants of all the enzymes used in this study (**Supplementary Table 4**) were transformed into chemically competent *E. coli* strains. 3081/211 were used to transform BspQI through electroporation. T7 Express *lysY*/Iq were used to transform T7 RNAP, FCE, FCE::T7 RNAP and 2'OMTase through heat shock. Single colonies of *E. coli* containing a plasmid encoding each protein were inoculated into 10 mL of Lysogeny broth (LB) medium containing: 30 µg mL$^{-1}$ of kanamycin and 10 µg mL$^{-1}$ of chloramphenicol for BspQI, 30 µg mL$^{-1}$ of ampicillin for T7 RNAP and 30 µg mL$^{-1}$ of kanamycin for FCE, FCE::T7 RNAP and 2'OMTase. Cells were grown for 2 hours at 37 °C with shaking at 220 rpm. The culture was diluted 1:100 into 1 L of LB medium with the corresponding antibiotics (at the same concentration), and the culture was grown at 37 °C until OD600nm reached 0.6-0.8. At that point, the protein expression was induced by adding isopropyl-D-thiogalactoside (IPTG) to a final concentration of 0.3 mM, and the cultures were incubated overnight at 16 °C with shaking at 220 rpm. Cells were collected by centrifugation (4000 rpm for 30 min at 4 °C) and cell pellet was resuspended with 50 mL of 20 mM Tris-HCl (pH 7.5), 50 mM NaCl, 20 mM imidazole, 20 mM L-Arg, 2 % glycerol. Cells were lysed by sonication on ice and cell debris was removed by centrifugation (13000 rpm for 30 min).

Proteins were purified through three different sequential columns. Ion-exchange diethylaminoethyl (HiTrap® DEAE) cellulose chromatography was used for the separation of proteins from nucleic acids. The column flow-through was recovered and loaded in HisTrap® Ni Sepharose for His-tag affinity purification. His-tagged proteins were eluted with a gradient, using as second buffer 20 mM Tris-HCl (pH 7.5), 150 mM NaCl, 500 mM imidazole, 20 mM L-Arg. Eluted fractions were combined and dialyzed in 20 mM Bis-Tris (pH 6.5), 50 mM NaCl, 1 mM DTT, 20 mM L-Arg. The combined fraction was loaded in HiTrap® Heparin column, and proteins were eluted in gradient using as second buffer 20 mM BisTris (pH 6.5), 1 M NaCl, 1 mM DTT, 20 mM L-Arg. Protein concentration was qualitatively determined for each fraction through sodium dodecyl sulfate polyacrylamide electrophoresis (SDS-PAGE, **Supplementary Fig.11**) and quantitatively determined through the Bradford protein assay. Fractions with the protein of interest were dialyzed in 40 mM Tris-HCl (pH 8.0), 100 mM NaCl, 50 mM L-Arg, 0.1 mM TCEP, 50 % glycerol, and stored at - 20 °C for short-term and at - 80 °C for long-term usage.

##### SNAP-tagged proteins immobilization in BG Magnetic Beads

BG- or BG-PEG-functionalized magnetic beads' supernatant was removed by using magnetic separation rack. Beads were washed five times with 137 mM NaCl, 2.6 mM KCl, 10 mM $Na_2HPO_4$, 1.76 mM $KH_2PO_4$, 0.005 % sodium azide (pH 8.0). SNAP-tagged protein of interest was diluted in the same buffer to a final concentration of 5 µM. The beads were resuspended with enzyme load (to a final bead concentration of 10 mg $mL^{-1}$) and incubated for 2 hours at room temperature. The flow-through was recovered and beads were washed again five times with the same buffer. Beads were resuspended in 40 mM Tris-HCl (pH 8.0), 100 mM NaCl, 50 mM L-Arg, 0.1 mM TCEP, 50 % glycerol (to a final bead concentration of 10 mg $mL^{-1}$) and stored at - 20 °C. Immobilization yield was qualitatively determined for each fraction through SDS-PAGE and quantitatively determined through the Bradford protein assay (**Supplementary Fig.1, 12**).

##### Calculation of immobilization parameters

The immobilization parameters characterized in this study were calculated as follows:

The immobilized protein is the mass of immobilized protein per gram of carrier.

$$\text{Immobilized protein (mg g}^{-1}\text{)} =$$

$$\left(\text{offered protein (mg mL}^{-1}\text{)} - \text{protein in supernatant (mg mL}^{-1}\text{)}\right) \text{ x } \frac{\text{immobilization volume (mL)}}{\text{carrier mass (g)}}$$

### Supplementary Figures

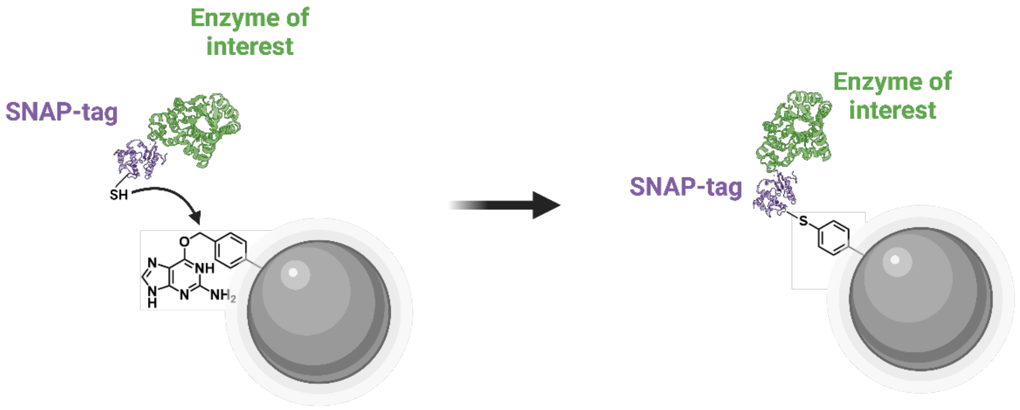


**Supplementary Figure 1.** Immobilization strategy for SNAP-tagged enzymes on benzyl guanine coated Sera-Mag™ microbeads.

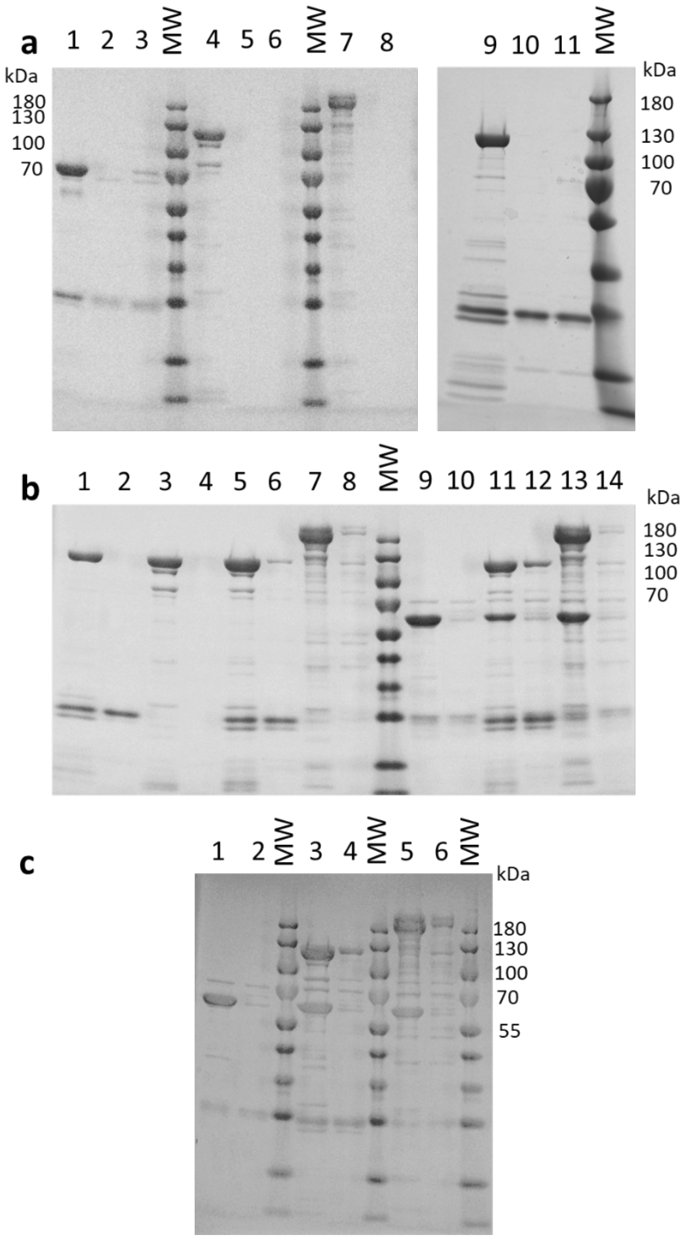


**Supplementary Figure 2.** Monitoring of enzyme immobilization by SDS-PAGE. MW: molecular weight marker. **(a)** Immobilization of enzymes on BG- and BG-$PEG_{750}$-coated magnetic beads. 1: BspQI load (~200 µg); 2: flow-through of magnetic BG beads BspQI immobilization; 3: flow-through of magnetic BG-$PEG_{750}$ beads BspQI immobilization; 4: FCE load (~200 µg); 5: flow-through of magnetic BG beads FCE immobilization; 6: flow-through of magnetic BG-$PEG_{750}$ beads FCE immobilization; 7: FCE::T7 RNAP load (~200 µg); 8: flow-through of magnetic BG-$PEG_{750}$ beads FCE::T7 RNAP immobilization; 9: T7 RNAP load (~200 µg). 10: flow-through of magnetic BG beads T7 RNAP immobilization. 11: flow-through of magnetic BG-$PEG_{750}$ beads T7 RNAP immobilization. **(b)** Immobilization and co-immobilization on BG-coated magnetic beads. 1: T7 RNAP load; 2: flow-through of magnetic BG-beads T7 RNAP immobilization; 3:

FCE load; 4: flow-through of magnetic BG-beads FCE immobilization; 5: T7 RNAP+FCE load; 6: flow-through of magnetic BG-beads T7 RNAP+FCE co-immobilization; 7: FCE::T7 RNAP load; 8: flow-through of magnetic BG-beads FCE::T7 RNAP immobilization; 9: 2'OMTase load; 10: flow-through of magnetic BG-beads 2'OMTase immobilization; 11: T7 RNAP+FCE+2'OMTase load; 12: flow-through of magnetic BG-beads T7 RNAP+FCE+2'OMTase co-immobilization; 13: FCE::T7 RNAP+2'OMTase load; 14: flow-through of magnetic BG-beads FCE::T7 RNAP+2'OMTase co-immobilization. **(c)** Immobilization and co-immobilization on BG-$PEG_{750}$-coated magnetic beads. 1: 2'OMTase load; 2: flow-through of magnetic BG-$PEG_{750}$-beads 2'OMTase immobilization; 3: T7 RNAP+FCE+2'OMTase load; 4: flow-through of magnetic BG-$PEG_{750}$-beads T7 RNAP+FCE+2'OMTase co-immobilization; 5: FCE::T7 RNAP+2'OMTase load; 6: flow-through of magnetic BG-$PEG_{750}$-beads FCE::T7 RNAP+2'OMTase co-immobilization.

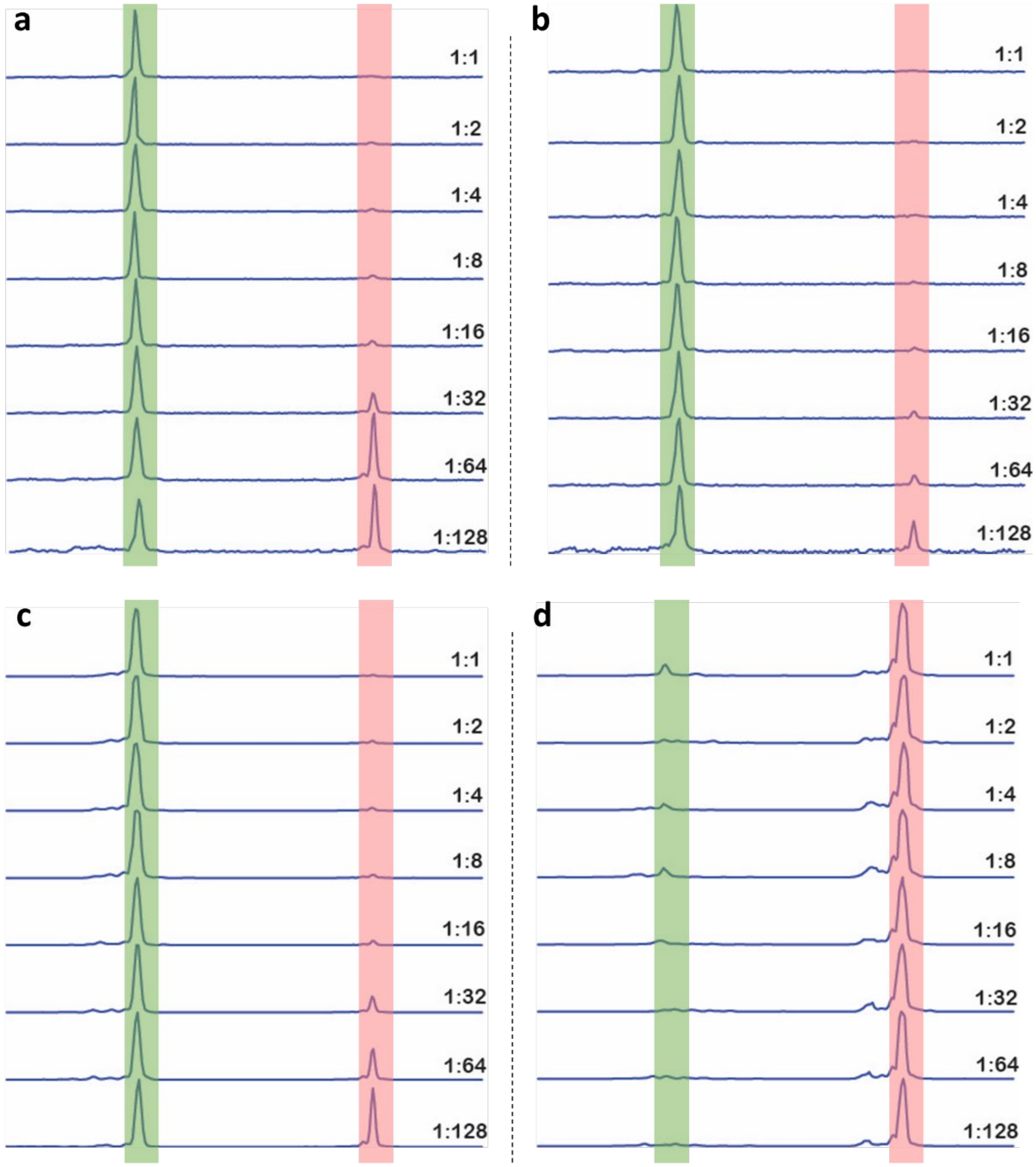


**Supplementary Figure 3.** Capillary electrophoresis traces of FAM-labeled 60-mer DNA substrate cleavage with BspQI at different dilutions (starting at 1 μM enzyme concentration). The major peak on the left (shadowed in green) indicates cleaved DNA, and the major peak on the right (shadowed in red) indicates the uncleaved DNA. **(a)** Free BspQI in NEB 3.1 buffer. **(b)** Free BspQI in RNA polymerase reaction buffer. **(c)** BspQI@BG in RNA polymerase reaction buffer. **(d)** Immobilization flow-through in RNA polymerase reaction buffer.

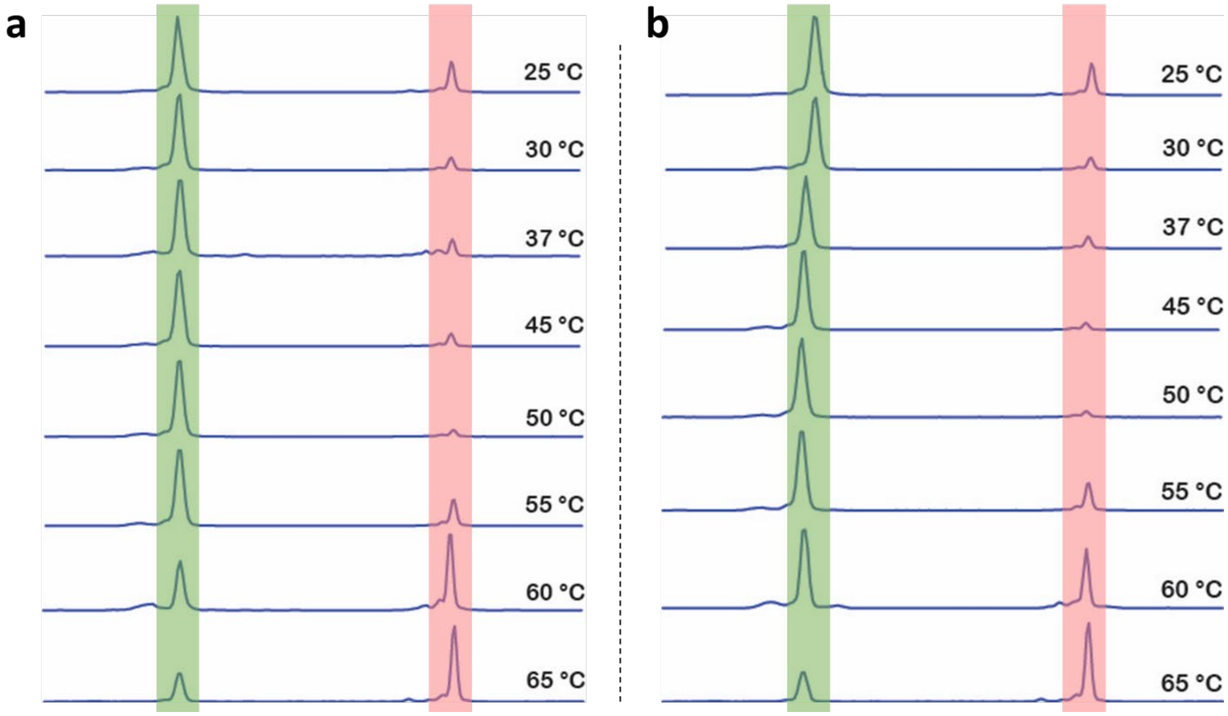


**Supplementary Figure 4.** Capillary electrophoresis traces of FAM-labeled 60-mer DNA substrate cleavage with BspQI (at 1 μM enzyme concentration) incubated at different temperatures (from 25 to 65 °C) for 1 h. The major peak on the left (shadowed in green) indicates cleaved DNA, and the major peak on the right (shadowed in red) indicates the uncleaved DNA. **(a)** Free BspQI. **(b)** BspQI@BG.

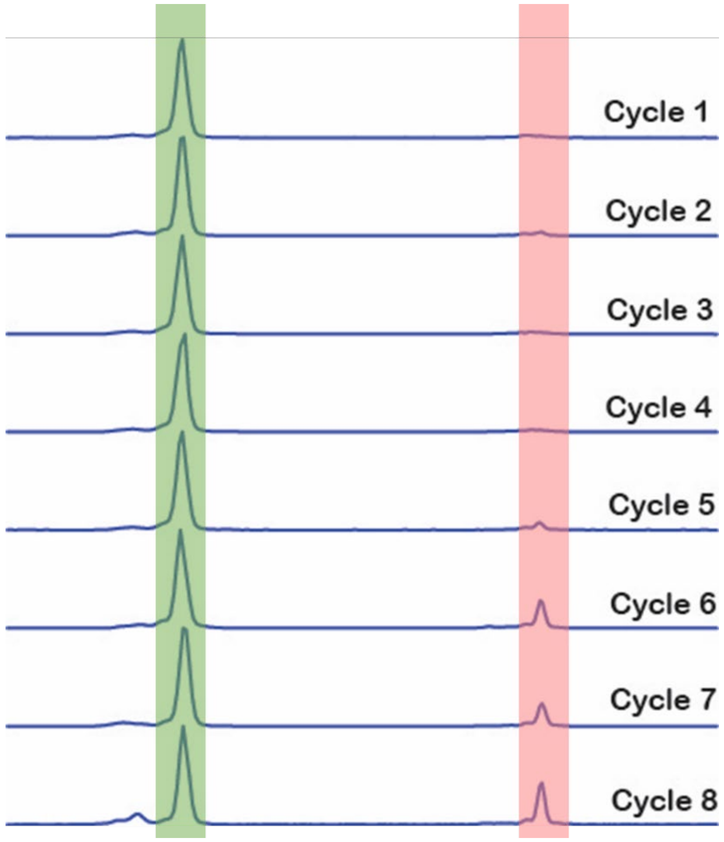


**Supplementary Figure 5.** Capillary electrophoresis traces of FAM-labeled 60-mer DNA substrate cleavage with BspQI@BG (at 1 µM of enzyme concentration) incubated at 50 °C for subsequent operation cycles of 1 h.

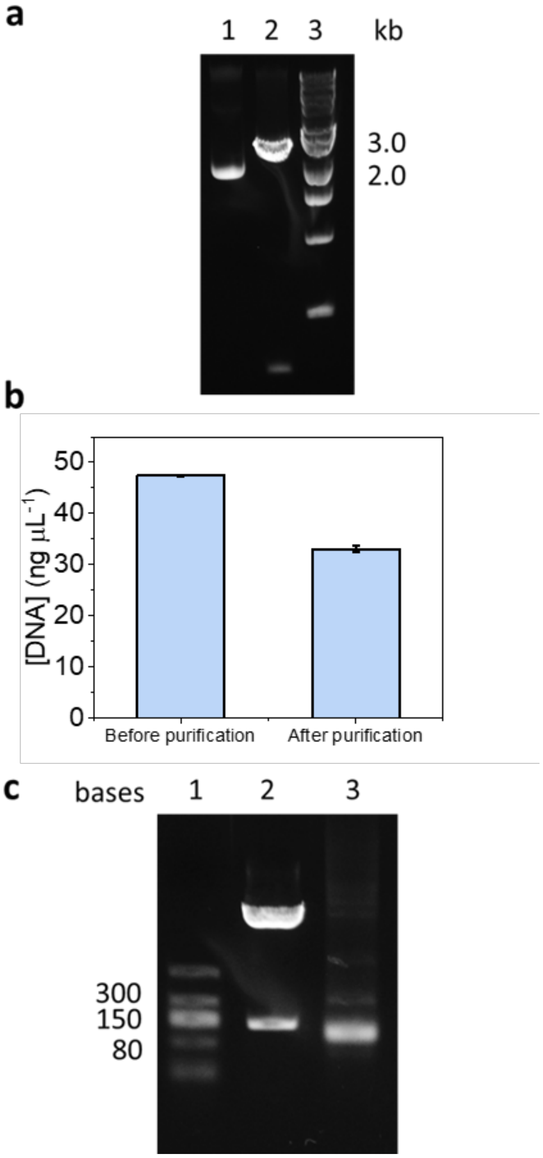


**Supplementary Figure 6. (a)** 1 % agarose gel showing pRNA21 linearization using BspQI@BG. Samples were incubated with Gel Loading Dye, Purple (6X) (NEB #B7024S). Lane 1: plasmid pRNA21 negative control; Lane 2: linearized pRNA21 DNA after incubation with 0.15 µM BspQI@BG at 50 °C for 1h; Lane 3: 1 kb DNA ladder (NEB # N3232S). **(b)** Linear pRNA21 DNA concentration before and after column purification. **(c)** 1 % agarose gel of pRNA21 *in vitro* transcription. Samples were incubated with RNA Loading Dye (2X) (NEB # B0363S). Lane 1: Low Range ssRNA Ladder (NEB # N0364S); Lane 2: Linearized pRNA21 plasmid after *in vitro* transcription with T7 RNA polymerase; Lane 3: Linearized pRNA21 plasmid *in vitro* transcription with T7 RNA polymerase after DNaseI digestion and RNA column purification.

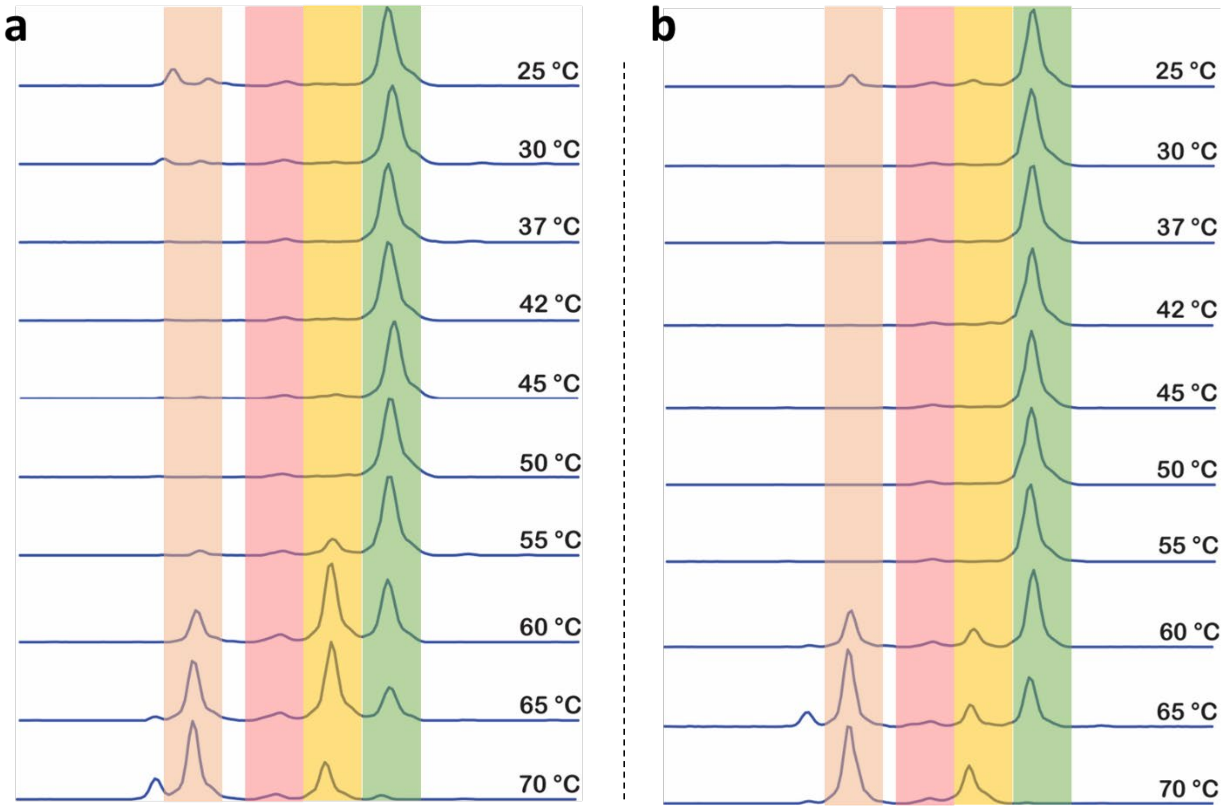


**Supplementary Figure 7.** Capillary electrophoresis traces of FCE capping efficiency using 3' FAM-labeled RNA oligonucleotide (ppp25mer) as substrate with FCE (at 0.2 µM of enzyme concentration) incubated at different temperatures (from 25 to 70 °C) for 30 min. The peak shadowed in red indicates the substrate (ppp-), the peak shadowed in orange indicates the product of the triphosphatase activity (pp-), the peak shadowed in yellow indicates the product of the guanylyl transferase activity (Gppp-) and the peak shadowed in green indicates the product of the N7-methyltransferase activity (m7Gppp-). **(a)** Free FCE. **(b)** FCE@PEG.

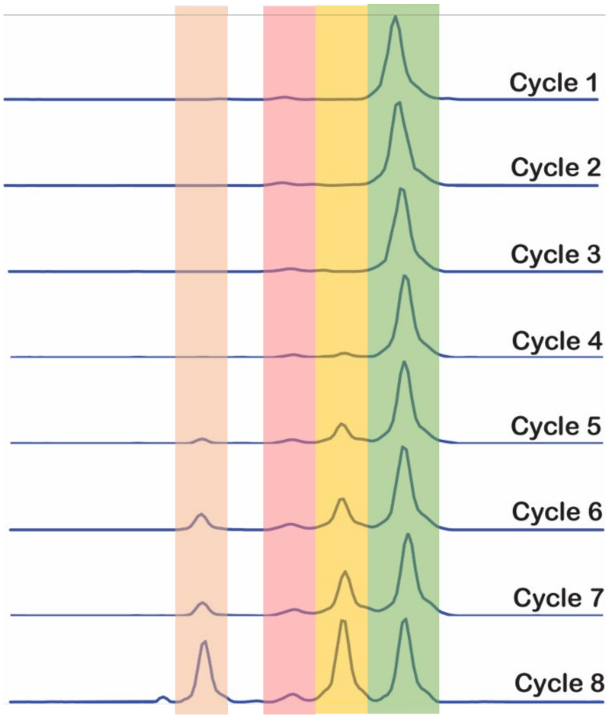


**Supplementary Figure 8.** Capillary electrophoresis traces of capping reaction with 3' FAM-labeled RNA oligonucleotide (ppp25mer) and FCE@PEG (at 0.2 µM enzyme concentration) incubated at 45 °C for subsequent operation cycles of 30 min.

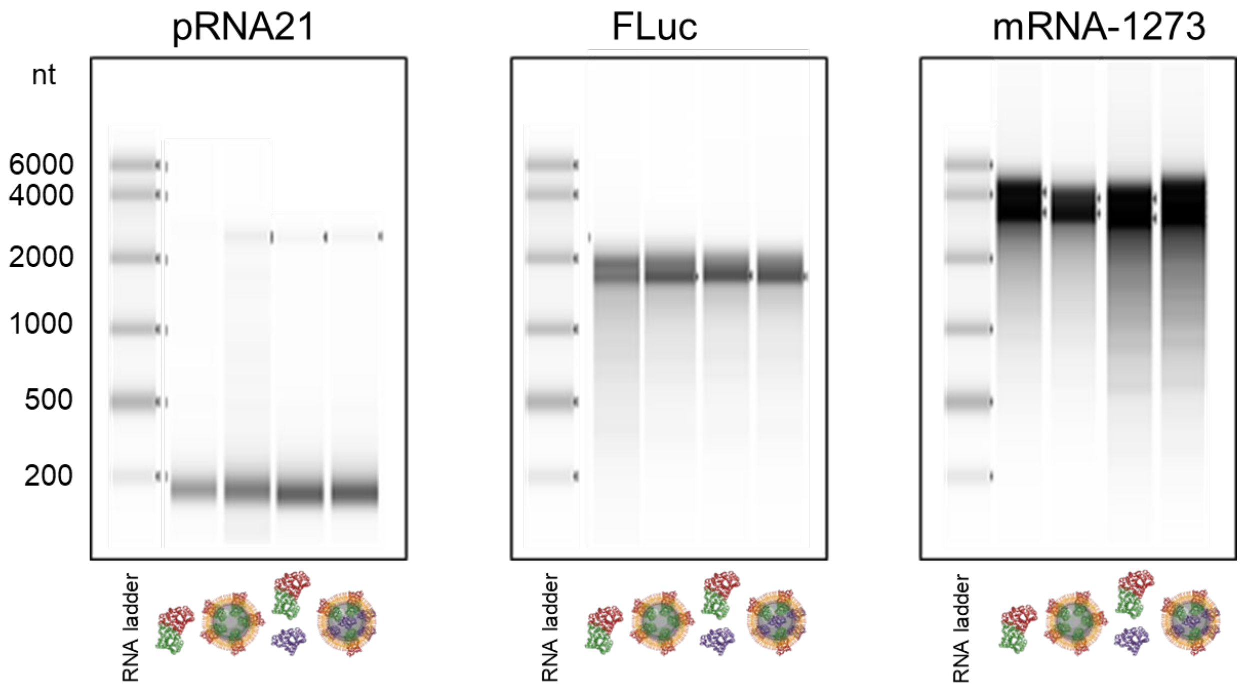


**Supplementary Figure 9.** Gel electrophoresis of co-transcriptional capping products by FCE::T7RNAP (in its soluble or immobilized form, and in presence of 2'-OMTase, both in solution and in co-immobilized form) after purification using Monarch RNA Clean-up kit (50 µg scale; New England Biolabs). pRNA21 (173 nt), FLuc (2118 nt) and mRNA-1273* (4146 nt) were analyzed on 2% E-gel (ThermoFisher). The bands were visualized using SYBR Gold and scanned using Amersham Typhoon RGB scanner with the Cy2 channel.

### Supplementary Tables

| Enzyme | Material | Chemical groups | Immobilized protein (mg $g^{-1}$ support) |
|---|---|---|---|
| BspQI | Sera-Mag™ Beads | BG | ~20 |
| | | BG, PEG750 | |
| T7 RNAP | | BG | |
| | | BG, PEG750 | |
| FCE | | BG | |
| | | BG, PEG750 | |
| FCE::T7 RNAP | | BG | |
| | | BG, PEG750 | |
| 2'OMTase | | BG | |
| | | BG, PEG750 | |

**Supplementary Table 1.** Estimated immobilization efficiency for different enzymes, based on Bradford assay, determining protein concentration in the load fraction and in the flow-through fraction.

| Target enzyme | Size (bp) | DNA/RNA sequence |
|---|---|---|
| BspQI | 60 | 5'/FAM/TGCCGCTTTCTGCATCAGCACATCATCTTCAGGCTCTTCGTCAGCCTCGCGCCG GTTCAG/3' |
| FCE/2'OMTase | 25 | 5'-ppp/GUAGAACUUCGUCGAGUACGCUCAA/3'-FAM |

**Supplementary Table 2.** Sequences of FAM-labeled DNA and RNA oligonucleotide substrates used for capillary electrophoresis and LCMS assays.

| DNA | Sequence |
|---|---|
| pRNA21 | 5’/TCGCGAATGCGTCGAGATTAATACGACTCACTATAGGGAAATAAGAGAGAAA<br>AGAAGAGTAAGAAGAAATATAAGAGCCACCATGGTTAACGCTGCCTTCTGCGG<br>GGCTTGCCTTCTGGCCATGCCCTTCTTCTCTCCCTTGCACCTGTACCTCTTGGTCT<br>TTGAATAAAGCCTGAGTAGGAAGAAAAAAAAAAAAAAAAAAAAAAAAA**GAAG**<br>**AGC**TAG/3’ |
| FLuc | 5’/GCTAGCCCCGCGAAATTAATACGACTCACTATAGGGTCTAGAAATAATTTTGT<br>TTAACTTTAAGAAGGAGATATAACCATGAAAATCGAAGAAGGTAAAGGTCACC<br>ATCACCATCACCACGGATCCATGGAAGACGCCAAAAACATAAAGAAAGGCCCG<br>GCGCCATTCTATCCTCTAGAGGATGGAACCGCTGGAGAGCAACTGCATAAGGCT<br>ATGAAGAGATACGCCCTGGTTCCTGGAACAATTGCTTTTACAGATGCACATATC<br>GAGGTGAACATCACGTACGCGGAATACTTCGAAATGTCCGTTCGGTTGGCAGAA<br>GCTATGAAACGATATGGGCTGAATACAAATCACAGAATCGTCGTATGCAGTGAA<br>AACTCTCTTCAATTCTTTATGCCGGTGTTGGGCGCGTTATTTATCGGAGTTGCAG<br>TTGCGCCCGCGAACGACATTTATAATGAACGTGAATTGCTCAACAGTATGAACA<br>TTTCGCAGCCTACCGTAGTGTTTGTTTCCAAAAAGGGGTTGCAAAAAATTTTGAA<br>CGTGCAAAAAAAATTACCAATAATCCAGAAAATTATTATCATGGATTCTAAAAC<br>GGATTACCAGGGATTTCAGTCGATGTACACGTTCGTCACATCTCATCTACCTCCC<br>GGTTTTAATGAATACGATTTTGTACCAGAGTCCTTTGATCGTGACAAAACAATTG<br>CACTGATAATGAATTCCTCTGGATCTACTGGGTTACCTAAGGGTGTGGCCCTTCC<br>GCATAGAACTGCCTGCGTCAGATTCTCGCATGCCAGAGATCCTATTTTTGGCAAT<br>CAAATCATTCCGGATACTGCGATTTTAAGTGTTGTTCCATTCCATCACGGTTTTG<br>GAATGTTTACTACACTCGGATATTTGATATGTGGATTTCGAGTCGTCTTAATGTA<br>TAGATTTGAAGAAGAGCTGTTTTTACGATCCCTTCAGGATTACAAAATTCAAAGT<br>GCGTTGCTAGTACCAACCCTATTTTCATTCTTCGCCAAAAGCACTCTGATTGACA<br>AATACGATTTATCTAATTTACACGAAATTGCTTCTGGGGGCGCACCTCTTTCGAA<br>AGAAGTCGGGGAAGCGGTTGCAAAACGCTTCCATCTTCCAGGGATACGACAAG<br>GATATGGGCTCACTGAGACTACATCAGCTATTCTGATTACACCCGAGGGGGATG<br>ATAAACCGGGCGCGGTCGGTAAAGTTGTTCCATTTTTTGAAGCGAAGGTTGTGG<br>ATCTGGATACCGGGAAAACGCTGGGCGTTAATCAGAGAGGCGAATTATGTGTCA<br>GAGGACCTATGATTATGTCCGGTTATGTAAACAATCCGGAAGCGACCAACGCCT<br>TGATTGACAAGGATGGATGGCTACATTCTGGAGACATAGCTTACTGGGACGAAG<br>ACGAACACTTCTTCATAGTTGACCGCTTGAAGTCTTTAATTAAATACAAAGGATA<br>TCAGGTGGCCCCCGCTGAATTGGAATCGATATTGTTACAACACCCCAACATCTTC<br>GACGCGGGCGTGGCAGGTCTTCCCGACGATGACGCCGGTGAACTTCCCGCCGCC<br>GTTGTTGTTTTGGAGCACGGAAAGACGATGACGGAAAAAGAGATCGTGGATTAC<br>GTCGCCAGTCAAGTAACAACCGCGAAAAAGTTGCGCGGAGGAGTTGTGTTTGTG<br>GACGAAGTACCGAAAGGTCTTACCGGAAAACTCGACGCAAGAAAAATCAGAGA<br>GATCCTCATAAAGGCCAAGAAGGGCGGAAAGTCCAAACTCGAGTAAG**GTTAAC**/<br>3’ |
| mRNA-1273* | 5’/CATATGGCTAGCCCCGCGAAATTAATACGACTCACTATAGGAAATAAGAGAG<br>AAAAGAAGAGTAAGAAGAAATATAAGACCCCGGCGCCGCCACCATGTTCGTGT<br>TCCTGGTGCTGCTGCCCCTGGTGAGCAGCCAGTGCGTGAACCTGACCACCCGGA<br>CCCAGCTGCCACCAGCCTACACCAACAGCTTCACCCGGGGCGTCTACTACCCCG<br>ACAAGGTGTTCCGGAGCAGCGTCCTGCACAGCACCCAGGACCTGTTCCTGCCCT<br>TCTTCAGCAACGTGACCTGGTTCCACGCCATCCACGTGAGCGGCACCAACGGCA<br>CCAAGCGGTTCGACAACCCCGTGCTGCCCTTCAACGACGGCGTGTACTTCGCCA<br>GCACCGAGAAGAGCAACATCATCCGGGGCTGGATCTTCGGCACCACCCTGGACA<br>GCAAGACCCAGAGCCTGCTGATCGTGAATAACGCCACCAACGTGGTGATCAAGG<br>TGTGCGAGTTCCAGTTCTGCAACGACCCCTTCCTGGGCGTGTACTACCACAAGA<br>ACAACAAGAGCTGGATGGAGAGCGAGTTCCGGGTGTACAGCAGCGCCAACAAC<br>TGCACCTTCGAGTACGTGAGCCAGCCCTTCCTGATGGACCTGGAGGGCAAGCAG<br>GGCAACTTCAAGAACCTGCGGGAGTTCGTGTTCAAGAACATCGACGGCTACTTC<br>AAGATCTACAGCAAGCACACCCCAATCAACCTGGTGCGGGATCTGCCCCAGGGC<br>TTCTCAGCCCTGGAGCCCCTGGTGGACCTGCCCATCGGCATCAACATCACCCGGT<br>TCCAGACCCTGCTGGCCCTGCACCGGAGCTACCTGACCCCAGGCGACAGCAGCA<br>GCGGGTGGACAGCAGGCGCGGCTGCTTACTACGTGGGCTACCTGCAGCCCCGGA<br>CCTTCCTGCTGAAGTACAACGAGAACGGCACCATCACCGACGCCGTGGACTGCG<br>CCCTGGACCCTCTGAGCGAGACCAAGTGCACCCTGAAGAGCTTCACCGTGGAGA<br>AGGGCATCTACCAGACCAGCAACTTCCGGGTGCAGCCCACCGAGAGCATCGTGC<br>GGTTCCCCAACATCACCAACCTGTGCCCCTTCGGCGAGGTGTTCAACGCCACCC<br>GGTTCGCCAGCGTGTACGCCTGGAACCGGAAGCGGATCAGCAACTGCGTGGCCG |

| | |
|---|---|
| | ACTACAGCGTGCTGTACAACAGCGCCAGCTTCAGCACCTTCAAGTGCTACGGCG<br>TGAGCCCCACCAAGCTGAACGACCTGTGCTTCACCAACGTGTACGCCGACAGCT<br>TCGTGATCCGTGGCGACGAGGTGCGGCAGATCGCACCCGGCCAGACAGGCAAG<br>ATCGCCGACTACAACTACAAGCTGCCCGACGACTTCACCGGCTGCGTGATCGCC<br>TGGAACAGCAACAACCTCGACAGCAAGGTGGGCGGCAACTACAACTACCTGTA<br>CCGGCTGTTCCGGAAGAGCAACCTGAAGCCCTTCGAGCGGGACATCAGCACCGA<br>GATCTACCAAGCCGGCTCCACCCCTTGCAACGGCGTGGAGGGCTTCAACTGCTA<br>CTTCCCTCTGCAGAGCTACGGCTTCCAGCCCACCAACGGCGTGGGCTACCAGCC<br>CTACCGGGTGGTGGTGCTGAGCTTCGAGCTGCTGCACGCCCCAGCCACCGTGTG<br>TGGCCCCAAGAAGAGCACCAACCTGGTGAAGAACAAGTGCGTGAACTTCAACTT<br>CAACGGCCTTACCGGCACCGGCGTGCTGACCGAGAGCAACAAGAAATTCCTGCC<br>CTTTCAGCAGTTCGGCCGGGACATCGCCGACACCACCGACGCTGTGCGGGATCC<br>CCAGACCCTGGAGATCCTGGACATCACCCCTTGCAGCTTCGGCGGCGTGAGCGT<br>GATCACCCCAGGCACCAACACCAGCAACCAGGTGGCCGTGCTGTACCAGGACGT<br>GAACTGCACCGAGGTGCCCGTGGCCATCCACGCCGACCAGCTGACACCCACCTG<br>GCGGGTCTACAGCACCGGCAGCAACGTGTTCCAGACCCGGGCCGGTTGCCTGAT<br>CGGCGCCGAGCACGTGAACAACAGCTACGAGTGCGACATCCCCATCGGCGCCG<br>GCATCTGTGCCAGCTACCAGACCCAGACCAATTCACCCCGGAGGGCAAGGAGCG<br>TGGCCAGCCAGAGCATCATCGCCTACACCATGAGCCTGGGCGCCGAGAACAGCG<br>TGGCCTACAGCAACAACAGCATCGCCATCCCCACCAACTTCACCATCAGCGTGA<br>CCACCGAGATTCTGCCCGTGAGCATGACCAAGACCAGCGTGGACTGCACCATGT<br>ACATCTGCGGCGACAGCACCGAGTGCAGCAACCTGCTGCTGCAGTACGGCAGCT<br>TCTGCACCCAGCTGAACCGGGCCCTGACCGGCATCGCCGTGGAGCAGGACAAGA<br>ACACCCAGGAGGTGTTCGCCCAGGTGAAGCAGATCTACAAGACCCCTCCCATCA<br>AGGACTTCGGCGGCTTCAACTTCAGCCAGATCCTGCCCGACCCCAGCAAGCCCA<br>GCAAGCGGAGCTTCATCGAGGACCTGCTGTTCAACAAGGTGACCCTAGCCGACG<br>CCGGCTTCATCAAGCAGTACGGCGACTGCCTCGGCGACATAGCCGCCCGGGACC<br>TGATCTGCGCCCAGAAGTTCAACGGCCTGACCGTGCTGCCTCCCCTGCTGACCG<br>ACGAGATGATCGCCCAGTACACCAGCGCCCTGTTAGCCGGAACCATCACCAGCG<br>GCTGGACTTTCGGCGCTGGAGCCGCTCTGCAGATCCCCTTCGCCATGCAGATGG<br>CCTACCGGTTCAACGGCATCGGCGTGACCCAGAACGTGCTGTACGAGAACCAGA<br>AGCTGATCGCCAACCAGTTCAACAGCGCCATCGGCAAGATCCAGGACAGCCTGA<br>GCAGCACCGCTAGCGCCCTGGGCAAGCTGCAGGACGTGGTGAACCAGAACGCC<br>CAGGCCCTGAACACCCTGGTGAAGCAGCTGAGCAGCAACTTCGGCGCCATCAGC<br>AGCGTGCTGAACGACATCCTGAGCCGGCTGGACCCTCCCGAGGCCGAGGTGCAG<br>ATCGACCGGCTGATCACTGGCCGGCTGCAGAGCCTGCAGACCTACGTGACCCAG<br>CAGCTGATCCGGGCCGCCGAGATTCGGGCCAGCGCCAACCTGGCCGCCACCAAG<br>ATGAGCGAGTGCGTGCTGGGCCAGAGCAAGCGGGTGGACTTCTGCGGCAAGGG<br>CTACCACCTGATGAGCTTTCCCCAGAGCGCACCCCACGGAGTGGTGTTCCTGCA<br>CGTGACCTACGTGCCCGCCCAGGAGAAGAACTTCACCACCGCCCCAGCCATCTG<br>CCACGACGGCAAGGCCCACTTTCCCCGGGAGGGCGTGTTCGTGAGCAACGGCAC<br>CCACTGGTTCGTGACCCAGCGGAACTTCTACGAGCCCCAGATCATCACCACCGA<br>CAACACCTTCGTGAGCGGCAACTGCGACGTGGTGATCGGCATCGTGAACAACAC<br>CGTGTACGATCCCCTGCAGCCCGAGCTGGACAGCTTCAAGGAGGAGCTGGACAA<br>GTACTTCAAGAATCACACCAGCCCCGACGTGGACCTGGGCGACATCAGCGGCAT<br>CAACGCCAGCGTGGTGAACATCCAGAAGGAGATCGATCGGCTGAACGAGGTGG<br>CCAAGAACCTGAACGAGAGCCTGATCGACCTGCAGGAGCTGGGCAAGTACGAG<br>CAGTACATCAAGTGGCCCTGGTACATCTGGCTGGGCTTCATCGCCGGCCTGATC<br>GCCATCGTGATGGTGACCATCATGCTGTGCTGCATGACCAGCTGCTGCAGCTGC<br>CTGAAGGGCTGTTGCAGCTGCGGCAGCTGCTGCAAGTTCGACGAGGACGACAGC<br>GAGCCCGTGCTGAAGGGCGTGAAGCTGCACTACACCTGATAATAGGCTGGAGCC<br>TCGGTGGCCTAGCTTCTTGCCCCTTGGGCCTCCCCCCAGCCCCTCCTCCCCTTCCT<br>GCACCCGTACCCCCGTGGTCTTTGAATAAAGTCTGAGTGGGCGGCAAAAAAAAA<br>AAAAAAAAAAAAAAAAAAAAAAAAAAAAAAAAAAAAAAAAAAAAAAAAAAAA<br>AAAAAAAAAAAAAAAAAAAAAAAAAAAAAAAAAAAAAAAAAA**GAGACG**ACAT<br>GT /3’ |

**Supplementary Table 3.** Sequences of DNA templates used for IVT (pRNA21, FLuc, pTsin CLuc and mRNA-1273* (Spike-Covid19 Moderna vaccine). Only the sequence corresponding to the RNA transcripts and neighboring sequences are shown. Promoter sequences are underlined, initiating nucleotides are colored in red, and restriction enzyme sites (BspQI for pRNA21, HpaI for FLuc, and BsmBI for mRNA-1273*) are in bold face.

| Enzyme | Specific Productivity (mg RNA mg enzyme$^{-1}$ min$^{-1}$) |
|---|---|
| Free T7 RNAP | 0.87 |
| T7@BG | 0.29 |
| T7@PEG | 0.37 |

**Supplementary Table 4.** Specific productivities of free and immobilized T7 RNA polymerase calculated based on linear phase of reaction time course (see **Fig.3a**). Specific productivity indicates the mg of RNA product obtained in the reaction per mg of enzyme per minute.

| Enzyme | Sequence |
|---|---|

| | |
|---|---|
| BspQI | 5’/DKDCEMKRTTLDSPLGKLELSGCEQGLHEIKLLGKGTSAADAVEVPAPAAVLGG<br>PEPLMQATAWLNAYFHQPEAIEEFPVPALHHPVFQQESFTRQVLWKLLKVVKFGEV<br>ISYQQLAALAGNPAATAAVKTALSGNPVPILIPCHRVVSSSGAVGGYEGGLAVKEW<br>LLAHEGHRLGKPGLGPAAIGAPGSGSGSPAGGSGSGSMRRLAKNSRNDSYLSNRDY<br>QEIVRENTTTISFPLKEKHTLTLTKKIGLNQTAGFGGWFFPDSPCLLTVTVLSSFGTK<br>VTSKTFSLSKDWNRVGLAWINEHSSDTMSIVLEFSDVEIVHTWGLTCDVFNVHELII<br>DAIEDQNKLIDVLNQEHLSPETYYLNHDSDTDLIENLESTEEIKIVNQSQKQISLKKCC<br>YCQRYMPVNILVRSNSSFHKHKSKKTGFQNECRACKKWRINNSFNPVRTKDQLHES<br>AVITREKKILLKEPEILQKIKNRNNGEGLKSIIWKKFDKKCFNCEKELTIEEVRLDHTR<br>PLAYLWPIDEHATCLCEKCNNTKHDMFPIDFYQGDEDKLRRLARITGLDYESLVKR<br>DVNEVELARIINNIEDFATNVEARTFRSIRNKVKEVRPDTDLFEILKSKNINLYNELQ<br>YELLTRKD/3’ |
| T7 RNAP | 5’/MKRTTLDSPLGKLELSGCEQGLHEIKLLGKGTSAADAVEVPAPAAVLGGPEPLM<br>QATAWLNAYFHQPEAIEEFPVPALHHPVFQQESFTRQVLWKLLKVVKFGEVISYQQ<br>LAALAGNPAATAAVKTALSGNPVPILIPCHRVVSSSGAVGGYEGGLAVKEWLLAHE<br>GHRLGKPGLGNTINIAKNDFSDIELAAIPFNTLADHYGERLAREQLALEHESYEMGE<br>ARFRKMFERQLKAGEVADNAAAKPLITTLLPKMIARINDWFEEVKAKRGKRPTAFQ<br>FLQEIKPEAVAYITIKTTLACLTSADNTTVQAVASAIGRAIEDEARFGRIRDLEAKHF<br>KKNVEEQLNKRVGHVYKKAFMQVVEADMLSKGLLGGEAWSSWHKEDSIHVGVR<br>CIEMLIESTGMVSLHRQNAGVVGQDSETIELAPEYAEAIATRAGALAGISPMFQPCV<br>VPPKPWTGITGGGYWANGRRPLALVRTHSKKALMRYEDVYMPEVYKAINIAQNTA<br>WKINKKVLAVANVITKWKHCPVEDIPAIEREELPMKPEDIDMNPEALTAWKRAAAA<br>VYRKDKARKSRRISLEFMLEQANKFANHKAIWFPYNMDWRGRVYAVSMFNPQGN<br>DMTKGLLTLAKGKPIGKEGYYWLKIHGANCAGVDKVPFPERIKFIEENHENIMACA<br>KSPLENTWWAEQDSPFCFLAFCFEYAGVQHHGLSYNCSLPLAFDGSCSGIQHFSAM<br>LRDEVGGRAVNLLPSETVQDIYGIVAKKVNEILQADAINGTDNEVVTVTDENTGEIS<br>EKVKLGTKALAGQWLAYGVTRSVTKRSVMTLAYGSKEFGFRQQVLEDTIQPAIDSG<br>KGLMFTQPNQAAGYMAKLIWESVSVTVVAAVEAMNWLKSAAKLLAAEVKDKKT<br>GEILRKRCAVHWVTPDGFPVWQEYKKPIQTRLNLMFLGQFRLQPTINTNKDSEIDAH<br>KQESGIAPNFVHSQDGSHLRKTVVWAHEKYGIESFALIHDSFGTIPADAANLFKAVR<br>ETMVDTYESCDVLADFYDQFADQLHESQL/3’ |
| FCE | 5’/MKIEEDKDCEMKRTTLDSPLGKLELSGCEQGLHRIIFLGKGTSAADAVEVPAPAA<br>VLGGPEPLMQATAWLNAYFHQPEAIEEFPVPALHHPVFQQESFTRQVLWKLLKVVK<br>FGEVISYSHLAALAGNPAATAAVKTALSGNPVPILIPCHRVVQGDLDVGGYEGGLA<br>VKEWLLAHEGHRLGKPGLGSATASAKRLQRCQDVNQVCEIYNSKGGIGELELRFDK<br>LPQNLFAGVFDKLKPDGEIQTTMRVSNRDGVAREITFGGGVKTNEIFVKKQNICVFD<br>VVDIFSYKVAVSTEETVVEKPTMETTAGVRFKIRLSVEDVVKDWRIDLTAVKTAEL<br>GKIAQHTASIVQRTFPDNLLKLTGAEVAKLAADSYELELEYTGKSPATNEKVNVAA<br>KYAVELLSSVRNANSTAAASFGESVSDLCRVAKIIHTHEYANVVCRTPSFKMLLPQV<br>VSLTKSSYYGGLYPPENLWLAGKTDGVRALVVCEDGVAKVITAESVDITHGVCSAT<br>TILDCELNVDAKILYVFDVIISNNTQVYTQPFSTRITTDISDIKIDGYKIEMKPFVKVV<br>KADEATFKSAYKAPHNEGLIMIEDGAAYAATKTYKWKPLSHNTIDFLIKACPKQLIN<br>VDPYKPRAGYKLWLLFTTISLDQQRELGIEFIPAWKILFTDINMFGSRVPIQFQPAINP<br>LAYVCYLPEDVNVNDGDIVEMRAVDGYDTIPKWELVRSRNDRKNEPGFYGNNYKI<br>ASDIYLNYIDVFHFEDLYKYNPGYFEKNKSDIYVAPNKYRRYLIKSLFGRYLRDAK<br>WVIDAAAGRGADLHLYKAECVEHLLAIDIDPTAISELVRRRNEITGYNKSHRGGRN<br>MHSHRGQSHCAKSTSLHALVADLRENPDVLIPKIIQSRPHERCYDAIVINFAIHYLCD<br>TDEHIRDFLITVSRLLAPNGVFIFTTMDGESIVKLLADHKVRPGEAWTIHTGDVNSPD<br>STVPKYSIRRLYDSDKLTKTGQQIEVLLPMSGEMKAEPLCNIKNIISMARKMGLDLV<br>ESANFSVLYEAYARDYPDIYARMTPDDKLYNDLHTYAVFKRKKGASATSHHHHHH<br>HHST/3’ |
| FCE::T7 RNAP | 5’/MKIEEHHHHHHHHGAGTKIEEDKDCEMKRTTLDSPLGKLELSGCEQGLHRIIFLG<br>KGTSAADAVEVPAPAAVLGGPEPLMQATAWLNAYFHQPEAIEEFPVPALHHPVFQQ<br>ESFTRQVLWKLLKVVKFGEVISYSHLAALAGNPAATAAVKTALSGNPVPILIPCHRV<br>VQGDLDVGGYEGGLAVKEWLLAHEGHRLGKPGLGGATAGAKRLQRCQDVNQVC<br>EIYNSKGGIGELELRFDKLPQNLFAGVFDKLKPDGEIQTTMRVSNRDGVAREITFGG<br>GVKTNEIFVKKQNICVFDVVDIFSYKVAVSTEETVVEKPTMETTAGVRFKIRLSVED<br>VVKDWRIDLTAVKTAELGKIAQHTASIVQRTFPDNLLKLTGAEVAKLAADSYELEL<br>EYTGKSPATNEKVNVAAKYAVELLSSVRNANSTAAASFGESVSDLCRVAKIIHTHE<br>YANVVCRTPSFKMLLPQVVSLTKSSYYGGLYPPENLWLAGKTDGVRALVVCEDGV<br>AKVITAESVDITHGVCSATTILDCELNVDAKILYVFDVIISNNTQVYTQPFSTRITTDIS<br>DIKIDGYKIEMKPFVKVVKADEATFKSAYKAPHNEGLIMIEDGAAYAATKTYKWKP |

| | |
|---|---|
| | LSHNTIDFLIKACPKQLINVDPYKPRAGYKLWLLFTTISLDQQRELGIEFIPAWKILFT<br>DINMFGSRVPIQFQPAINPLAYVCYLPEDVNVNDGDIVEMRAVDGYDTIPKWELVRS<br>RNDRKNEPGFYGNNYKIASDIYLNYIDVFHFEDLYKYNPGYFEKNKSDIYVAPNKY<br>RRYLIKSLFGRYLRDAKWVIDAAAGRGADLHLYKAECVEHLLAIDIDPTAISELVRR<br>RNEITGYNKSHRGGRNMHSHRGQSHCAKSTSLHALVADLRENPDVLIPKIIQSRPHE<br>RCYDAIVINFAIHYLCDTDEHIRDFLITVSRLLAPNGVFIFTTMDGESIVKLLADHKVR<br>PGEAWTIHTGDVNSPDSTVPKYSIRRLYDSDKLTKTGQQIEVLLPMSGEMKAEPLCN<br>IKNIISMARKMGLDLVESANFSVLYEAYARDYPDIYARMTPDDKLYNDLHTYAVFK<br>RKKGAATAGTNTINIAKNDFSDIELAAIPFNTLADHYGERLAREQLALEHESYEMGE<br>ARFRKMFERQLKAGEVADNAAAKPLITTLLPKMIARINDWFEEVKAKRGKRPTAFQ<br>FLQEIKPEAVAYITIKTTLACLTSADNTTVQAVASAIGRAIEDEARFGRIRDLEAKHF<br>KKNVEEQLNKRVGHVYKKAFMQVVEADMLSKGLLGGEAWSSWHKEDSIHVGVR<br>CIEMLIESTGMVSLHRQNAGVVGQDSETIELAPEYAEAIATRAGALAGISPMFQPCV<br>VPPKPWTGITGGGYWANGRRPLALVRTHSKKALMRYEDVYMPEVYKAINIAQNTA<br>WKINKKVLAVANVITKWKHCPVEDIPAIEREELPMKPEDIDMNPEALTAWKRAAAA<br>VYRKDKARKSRRISLEFMLEQANKFANHKAIWFPYNMDWRGRVYAVSMFNPQGN<br>DMTKGLLTLAKGKPIGKEGYYWLKIHGANCAGVDKVPFPERIKFIEENHENIMACA<br>KSPLENTWWAEQDSPFCFLAFCFEYAGVQHHGLSYNCSLPLAFDGSCSGIQHFSAM<br>LRDEVGGRAVNLLPSETVQDIYGIVAKKVNEILQADAINGTDNEVVTVTDENTGEIS<br>EKVKLGTKALAGQWLAYGVTRSVTKRSVMTLAYGSKEFGFRQQVLEDTIQPAIDSG<br>KGLMFTQPNQAAGYMAKLIWESVSVTVVAAVEAMNWLKSAAKLLAAEVKDKKT<br>GEILRKRCAVHWVTPDGFPVWQEYKKPIQTRLNLMFLGQFRLQPTINTNKDSEIDAH<br>KQESGIAPNFVHSQDGSHLRKTVVWAHEKYGIESFALIHDSFGTIPADAANLFKAVR<br>ETMVDTYESCDVLADFYDQFADQLHESQLDKMPALPAKGNLNLRDILESDFAFA/3’ |
| 2’OMTase | 5’/HHHHHHGDKDCEMKRTTLDSPLGKLELSGCEQGLHEIKLLGKGTSAADAVEVPA<br>PAAVLGGPEPLMQATAWLNAYFHQPEAIEEFPVPALHHPVFQQESFTRQVLWKLLK<br>VVKFGEVISYQQLAALAGNPAATAAVKTALSGNPVPILIPCHRVVSSSGAVGGYEG<br>GLAVKEWLLAHEGHRLGKPGLGPAAIGAPGSGSGSPAGGSGSGSDVVSLDKPFMYF<br>EEIDNELDYEPESANEVAKKLPYQGQLKLLLGELFFLSKLQRHGILDGATVVYIGSA<br>PGTHIRYLRDHFYNLGVIIKWMLIDGRHHDPILNGLRDVTLVTRFVDEEYLRSIKKQ<br>LHPSKIILISDVRSKRGGNEPSTADLLSNYALQNVMISILNPVASSLKWRCPFPDQWI<br>KDFYIPHGNKMLQPFAPSYSAEMRLLSIYTGENMRLTRVTKSDAVNYEKKMYYLN<br>KIVRNKVVVNFDYPNQEYDYFHMYFMLRTVYCNKTFPTTKAKVLFLQQSIFRFLNIP<br>TTSTEKVSHEPIQRKISSKNSMSKNRNSKRSVRSNKLE/3’ |

**Supplementary Table 4.** Protein amino acid sequences for all the studied fusion enzymes. SNAP sequence is colored in yellow, BspQI sequence is colored in blue, T7 RNAP sequence is colored in red, FCE sequence is colored in green and 2’OMTase sequence is colored in purple.
